# From bud formation to flowering: transcriptomic state defines the cherry developmental phases of sweet cherry bud dormancy

**DOI:** 10.1101/586651

**Authors:** Noémie Vimont, Mathieu Fouché, José Antonio Campoy, Meixuezi Tong, Mustapha Arkoun, Jean-Claude Yvin, Philip A. Wigge, Elisabeth Dirlewanger, Sandra Cortijo, Bénédicte Wenden

## Abstract

**Background:** Bud dormancy is a crucial stage in perennial trees and allows survival over winter to ensure optimal flowering and fruit production. Recent work highlighted physiological and molecular events occurring during bud dormancy in trees. However, they usually examined bud development or bud dormancy in isolation. In this work, we aimed to further explore the global transcriptional changes happening throughout bud development and dormancy onset, progression and release.

**Results:** Using next-generation sequencing and modelling, we conducted an in-depth transcriptomic analysis for all stages of flower buds in several sweet cherry (*Prunus avium* L.) cultivars that are characterized for their contrasted dates of dormancy release. We find that buds in organogenesis, paradormancy, endodormancy and ecodormancy stages are defined by the expression of genes involved in specific pathways, and these are conserved between different sweet cherry cultivars. In particular, we found that *DORMANCY ASSOCIATED MADS-box (DAM)*, floral identity and organogenesis genes are up-regulated during the pre-dormancy stages while endodormancy is characterized by a complex array of signalling pathways, including cold response genes, ABA and oxidation-reduction processes. After dormancy release, genes associated with global cell activity, division and differentiation are activated during ecodormancy and growth resumption. We then went a step beyond the global transcriptomic analysis and we developed a model based on the transcriptional profiles of just seven genes to accurately predict the main bud dormancy stages.

**Conclusions:** Overall, this study has allowed us to better understand the transcriptional changes occurring throughout the different phases of flower bud development, from bud formation in the summer to flowering in the following spring. Our work sets the stage for the development of fast and cost effective diagnostic tools to molecularly define the dormancy stages. Such integrative approaches will therefore be extremely useful for a better comprehension of complex phenological processes in many species.

## BACKGROUND

Temperate trees face a wide range of environmental conditions including highly contrasted seasonal changes. Among the strategies to enhance survival under unfavourable climatic conditions, bud dormancy is crucial for perennial plants since its progression over winter is determinant for optimal growth, flowering and fruit production during the subsequent season. Bud dormancy has long been compared to an unresponsive physiological phase, in which metabolic processes within the buds are halted by cold temperature and/or short photoperiod. However, several studies have shown that bud dormancy progression can be affected in a complex way by temperature, photoperiod or both, depending on the tree species [1–5]. Bud dormancy has traditionally been separated into three main phases: (i) paradormancy, also named “summer dormancy” [6]; (ii) endodormancy, mostly triggered by internal factors; and (iii) ecodormancy, controlled by external factors [7, 8]. Progression through endodormancy requires cold accumulation whereas warmer temperatures, i.e. heat accumulation, drive the competence to resume growth over the ecodormancy phase. Dormancy is thus highly dependent on external temperatures, and changes in seasonal timing of bud break and blooming have been reported in relation with global warming. Notably, advances in bud break and blooming dates in spring have been observed for tree species, such as apple, cherry, birch, oak or Norway spruce, in the northern hemisphere, thus increasing the risk of late frost damages [9–14] while insufficient cold accumulation during winter may lead to incomplete dormancy release associated with bud break delay and low bud break rate [15, 16]. These phenological changes directly impact the production of fruit crops, leading to large potential economic losses [17]. Consequently, it becomes urgent to acquire a better understanding of bud responses to temperature stimuli in the context of climate change in order to tackle fruit losses and anticipate future production changes.

In the recent years, an increasing number of studies have investigated the physiological and molecular mechanisms of bud dormancy transitions in perennials using RNA sequencing technology, thereby giving a new insight into potential pathways involved in dormancy. The results suggest that the transitions between the three main bud dormancy phases (para-, endo- and eco- dormancy) are mediated by pathways related to *DORMANCY ASSOCIATED MADS-box (DAM)* genes [18], phytohormones [19–22], carbohydrates [22, 23], temperature [24, 25], photoperiod [26], reactive oxygen species [27, 28], water deprivation [26], cold acclimation and epigenetic regulation [29]. Owing to these studies, a better understanding of bud dormancy has been established in different perennial species [18, 30, 31]. However, we are still missing a fine-resolution temporal understanding of transcriptomic changes happening over the entire bud development, from bud organogenesis to bud break.

Indeed, the small number of sampling dates in existing studies seems to be insufficient to capture all the information about changes occurring throughout the dormancy cycle as it most likely corresponds to a chain of biological events rather than an on/off mechanism. Many unresolved questions remain: What are the fine-resolution dynamics of gene expression related to dormancy? Are specific sets of genes associated with dormancy stages? Since the timing for the response to environmental cues is cultivar-dependant [32, 33], are transcriptomic profiles during dormancy different in cultivars with contrasted flowering date?

To explore these mechanisms, we conducted a transcriptomic analysis of sweet cherry (*Prunus avium* L.) flower buds from bud organogenesis until the end of bud dormancy using next-generation sequencing. Sweet cherry is a perennial species highly sensitive to temperature [34] and we focused on three sweet cherry cultivars displaying contrasted flowering dates and response to environmental conditions. We carried out a fine-resolution time-course spanning the entire bud development, from flower organogenesis in July to flowering in spring of the following year (February to April), encompassing para-, enco- and ecodormancy phases. Our results indicate that transcriptional changes happening during dormancy are conserved between different sweet cherry cultivars, opening the way to the identification of key factors involved in the progression through bud dormancy.

## RESULTS

### Transcriptome accurately captures the dormancy state

In order to define transcriptional changes happening over the sweet cherry flower bud development, we performed a transcriptomic-wide analysis using next-generation sequencing from bud organogenesis to flowering. According to bud break percentage (Fig. 1a), morphological observations (Fig. 1b), average temperatures (see Fig. S1 in Additional File 1) and descriptions from Lang et al., (1987), we assigned five main stages to the flower buds samples (Fig. 1c): i) flower bud organogenesis occurs in July and August; ii) paradormancy corresponds to the period of growth cessation in September; iii) during the endodormancy phase, initiated in October, buds are unresponsive to forcing conditions therefore the increasing bud break percentage under forcing conditions suggests that endodormancy was released on 9^th^ December 2015, 29^th^ January 2016, and 26^th^ February 2016 for the three cultivars ‘Cristobalina’, ‘Garnet’ and ‘Regina’, respectively, thus corresponding to iv) dormancy breaking; and v) ecodormancy starting from the estimated dormancy release date until flowering. We harvested buds at eleven dates spanning all these bud stages for the sweet cherry cultivars ‘Cristobalina’, ‘Garnet’ and ‘Regina’, and generated a total of 81 transcriptomes (Table S1 in Additional File 2). First, in order to explore the transcriptomic characteristics of each bud stage separately from the cultivar effect, we focused the analysis on the early flowering cultivar ‘Garnet’.

**Figure 1.**
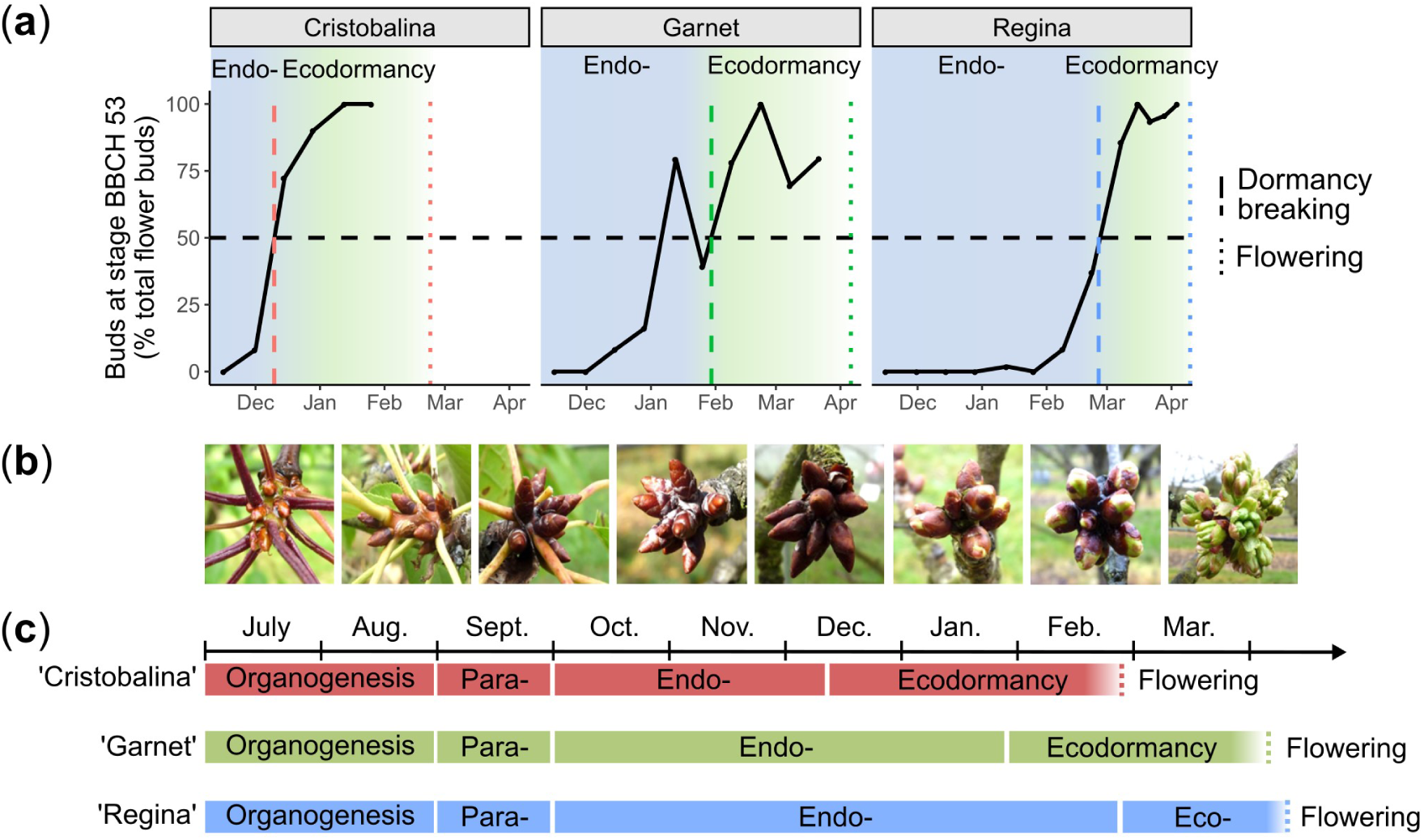
Dormancy status under environmental conditions and RNA-seq sampling dates. (a) Evaluation of bud break percentage under forcing conditions was carried out for three sweet cherry cultivars displaying different flowering dates in ‘Cristobalina’, ‘Garnet’ and ‘Regina’ for the early, medium and late cultivar, respectively. The dash and dotted lines correspond to the dormancy release date, estimated at 50% of buds at BBCH stage 53 [90], and the flowering date, respectively. (b) Pictures of the sweet cherry buds corresponding to the different sampling dates. (c) Sampling time points for the transcriptomic analysis are represented by coloured stars. Red for ‘Cristobalina, green for ‘Garnet’ and blue for ‘Regina’.

**Table 1.**
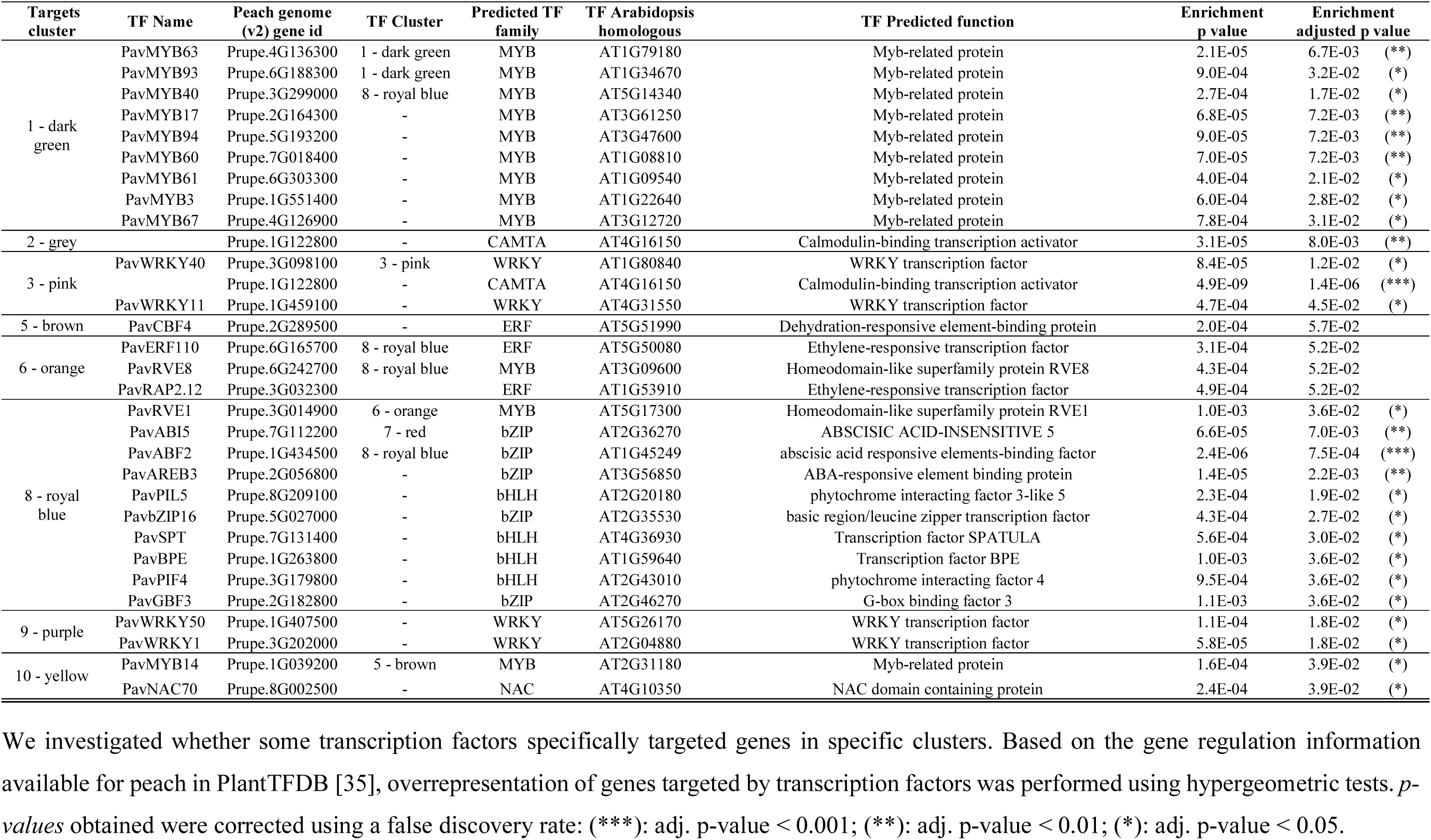
Transcription factors with over-represented targets in the different clusters

Using DESeq2 and a threshold of 0.05 on the adjusted *p-value*, we identified 6,683 genes that are differentially expressed (DEGs) between the defined bud stages for the sweet cherry cultivar ‘Garnet’ (Table S2 in Additional File 2). When projected into a two-dimensional space (Principal Component Analysis, PCA), data for these DEGs show that transcriptomes of samples harvested at a given date are projected together (Fig. 2), showing the high quality of the biological replicates and that different trees are in a very similar transcriptional state at the same date. Very interestingly, we also observe that flower bud states are clearly separated on the PCA, with the exception of organogenesis and paradormancy, which are projected together (Fig. 2). The first dimension of the analysis (PC1) explains 41,63% of the variance and clearly represents the strength of bud dormancy where samples on the right of the axis are in late endodormancy (Dec) or dormancy breaking stages, while samples on the left of the axis are in organogenesis and paradormancy. Samples harvested at the beginning of the endodormancy (Oct and Nov) are mid-way between samples in paradormancy and in late endodormancy (Dec) on PC1. The second dimension of the analysis (PC2) explains 20.24% of the variance and distinguishes two main phases of the bud development: before and after dormancy breaking. We obtain very similar results when performing the PCA on all genes (Fig. S2 in Additional File 1). These results indicate that the transcriptional state of DEGs accurately captures the dormancy state of flower buds.

**Figure 2.**
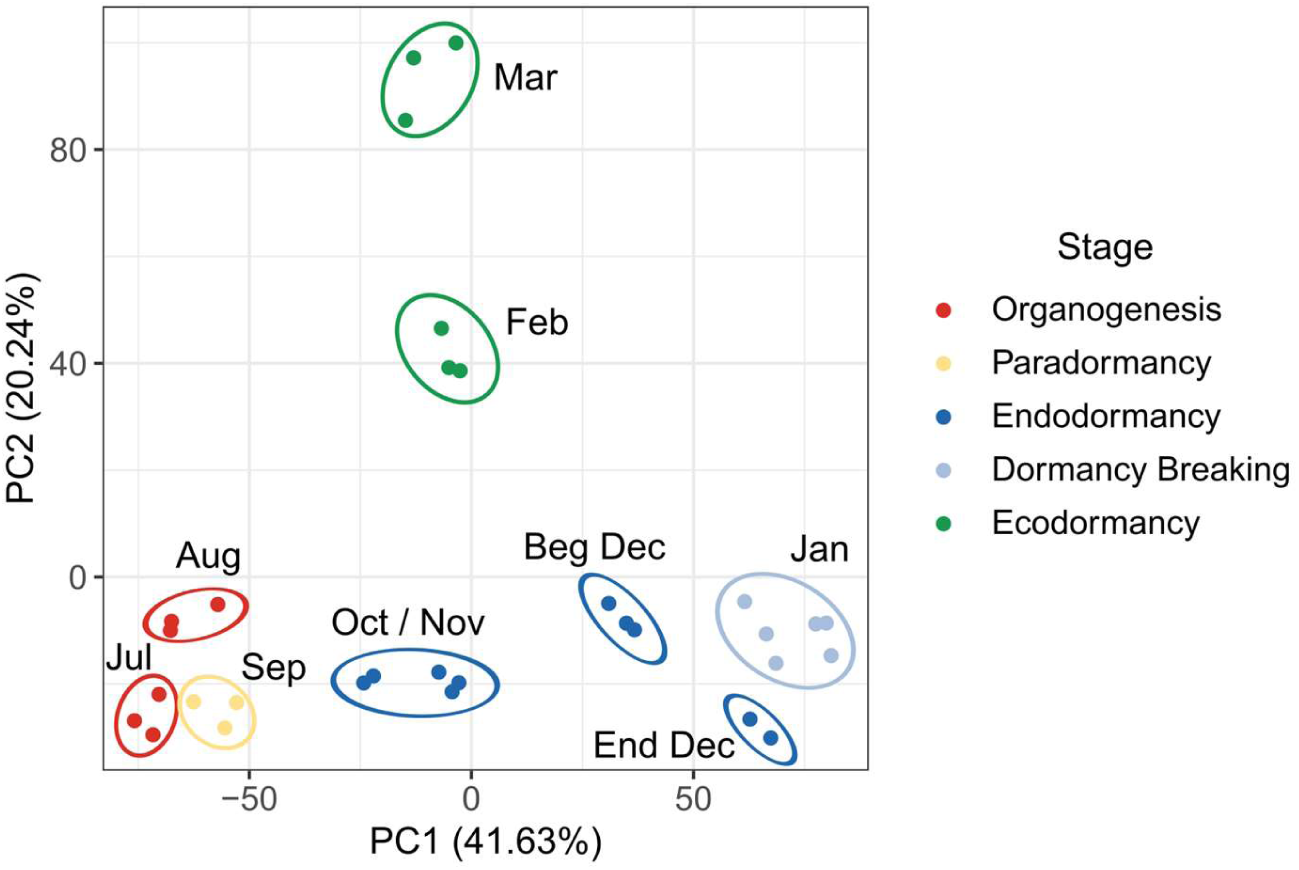
Separation of samples by dormancy stage using differentially expressed genes. The principal component analysis was conducted on the TPM (transcripts per millions reads) values for the differentially expressed genes in the cultivar ‘Garnet’ flower buds, sampled on three trees between July and March.

### Bud stage-dependent transcriptional activation and repression are associated with different pathways

We further investigated whether specific genes or signalling pathways could be associated with the different flower bud stages. For this, we performed a hierarchical clustering of the DEGs based on their expression in all samples. We could group the genes in ten clusters clearly showing distinct expression profiles throughout the bud development (Fig. 3). Overall, three main types of clusters can be discriminated: the ones with a maximum expression level during organogenesis and paradormancy (cluster 1: 1,549 genes; cluster 2: 70 genes; cluster 3: 113 genes; cluster 4: 884 genes and cluster 10: 739 genes, Fig. 3), the clusters with a maximum expression level during endodormancy and around the time of dormancy breaking (cluster 5: 156 genes; cluster 6: 989 genes; cluster 7: 648 genes and cluster 8: 612 genes, Fig. 3), and the clusters with a maximum expression level during ecodormancy (cluster 9: 924 genes and cluster 10: 739 genes, Fig. 3). This result shows that different groups of genes are associated with these three main flower bud phases. Interestingly, we also observed that during the endodormancy phase, some genes are expressed in October and November then repressed in December (cluster 4, Fig. 3), whereas another group of genes is expressed in December (clusters 8, 5, 6 and 7, Fig. 3) therefore separating endodormancy in two periods with distinct transcriptional states, which supports the PCA observation.

**Figure 3.**
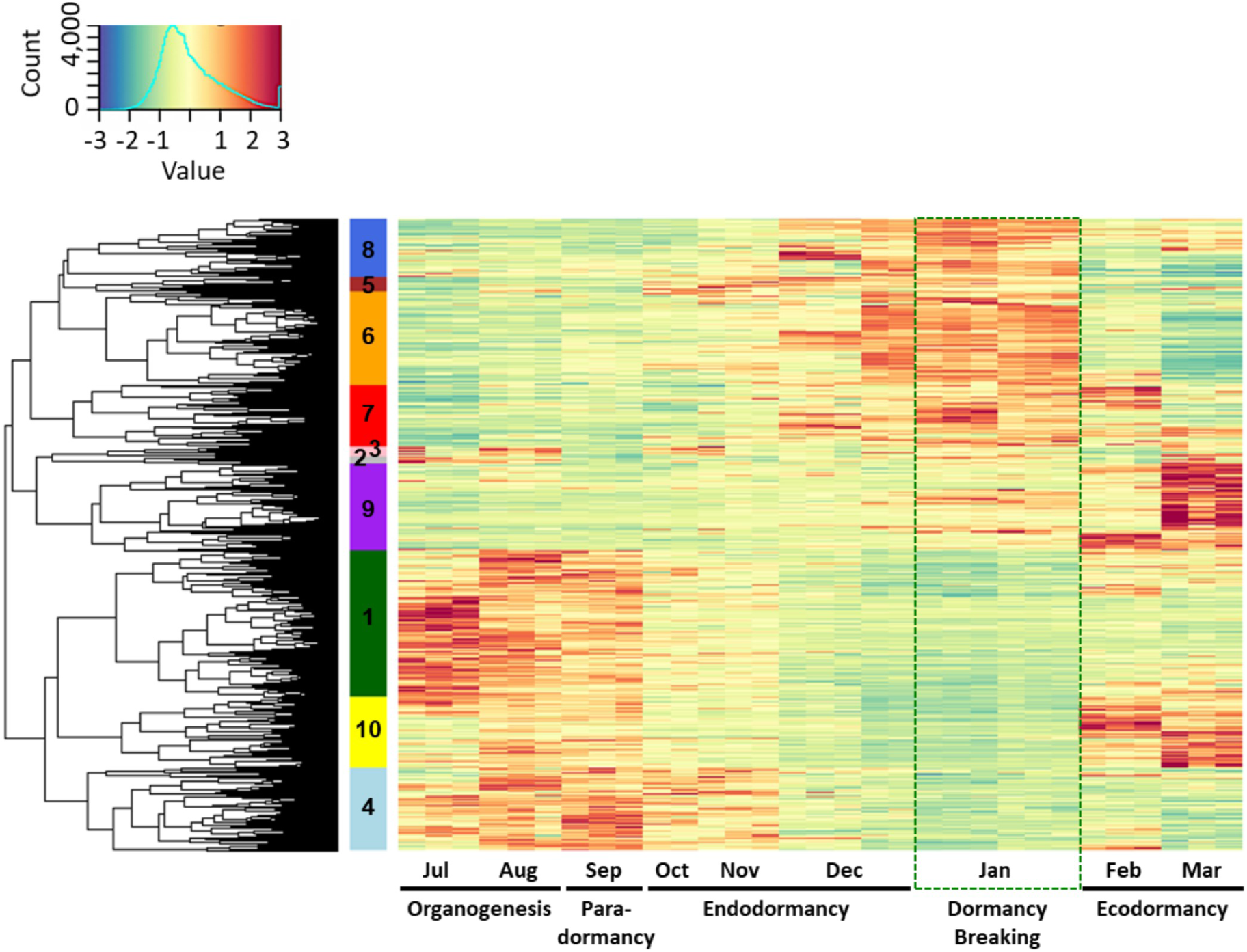
Clusters of expression patterns for differentially expressed genes in the sweet cherry cultivar ‘Garnet’. Heatmap for ‘Garnet’ differentially expressed genes during bud development. Each column corresponds to the gene expression for flower buds from one single tree at a given date. Clusters are ordered based on the chronology of the expression peak (from earliest – July, 1-dark green cluster – to latest – March, 9 and 10). Expression values were normalized and *z-scores* are represented here.

In order to explore the functions and pathways associated with the gene clusters, we performed a GO enrichment analysis for each of the ten identified clusters (Fig. 4, Fig. S3). GO terms associated with the response to stress as well as biotic and abiotic stimuli were enriched in the clusters 2, 3 and 4, with genes mainly expressed during organogenesis and paradormancy. In addition, we observed high expression of genes associated with floral identity before dormancy, including *AGAMOUS-LIKE20 (PavAGL20)* and the bZIP transcription factor *PavFD* (Fig. 5). On the opposite, at the end of the endodormancy phase (cluster 6, 7 and 8), we highlighted different enrichments in GO terms linked to basic metabolisms such as nucleic acid metabolic processes or DNA replication but also to response to alcohol and abscisic acid (ABA). For example, *ABA BINDING FACTOR 2 (PavABF2), ARABIDOPSIS THALIANA HOMEOBOX 7 (PavATHB7)* and ABA 8’-hydroxylase (*PavCYP707A2*), associated with the ABA pathway, as well as the stress-induced gene *PavHVA22*, were highly expressed during endodormancy (Fig. 5). During ecodormancy, genes in cluster 9 and 10 are enriched in functions associated with transport, cell wall biogenesis as well as oxidation-reduction processes (Fig. 4 and see Additional file 3). Indeed, we identified the *GLUTATHION S-TRANSFERASE8 (PavGST8)* gene and a peroxidase specifically activated during ecodormancy (Fig. 5). However, oxidation-reduction processes are likely to occur during endodormancy as well, as suggested by the expression patterns of *GLUTATHION PEROXIDASE 6 (PavGPX6*) and *GLUTATHION REDUCTASE (PavGR)*. Interestingly, *AGAMOUS (PavAG)* and *APETALA3 (PavAP3)* showed an expression peak during ecodormancy (Fig. 5). These results show that different functions and pathways are specific to flower bud development stages.

**Figure 4.**
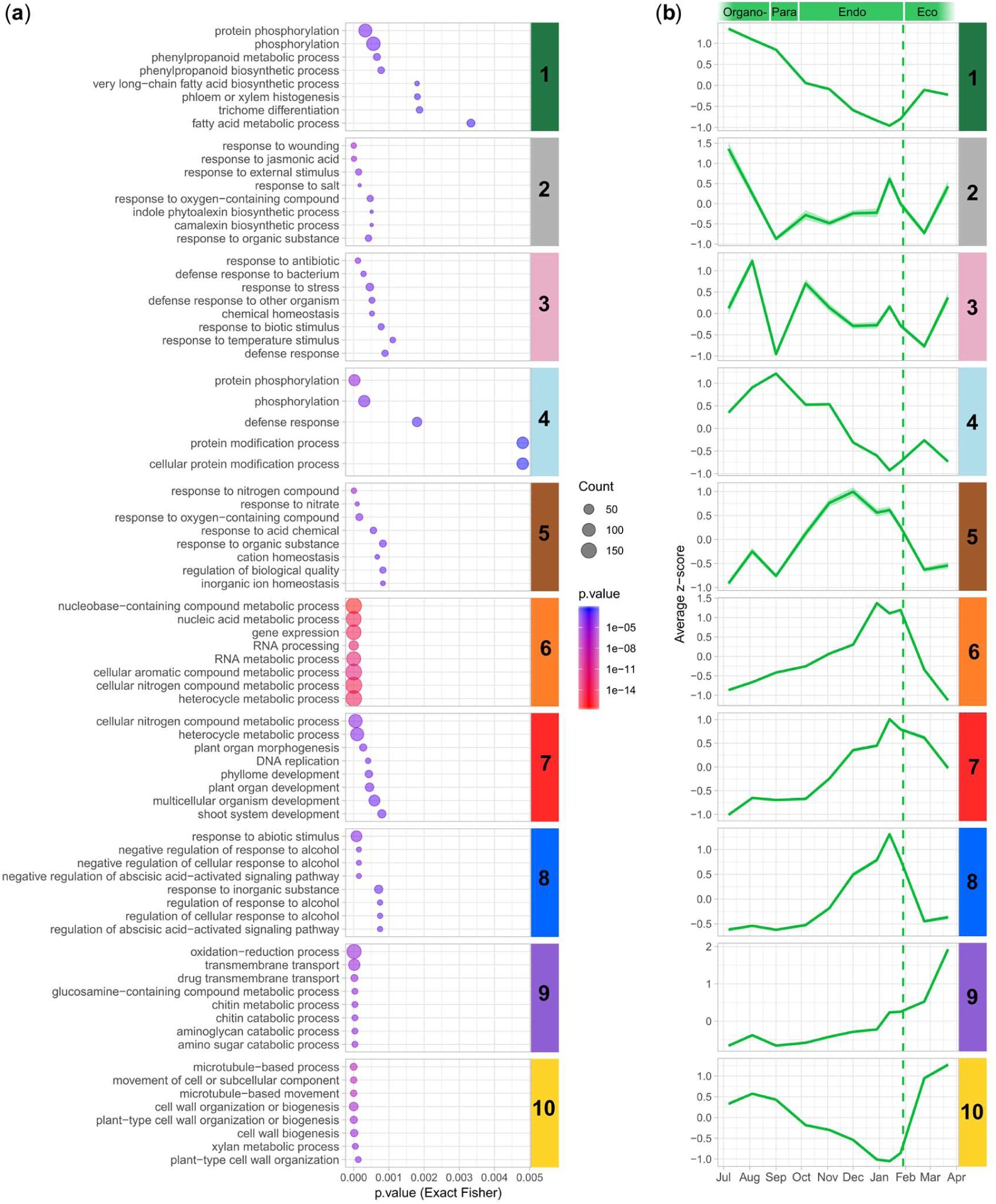
Enrichments in gene ontology terms for biological processes and average expression patterns in the different clusters in the sweet cherry cultivar ‘Garnet’. (a) Using the topGO package (Alexa & Rahnenführer, 2018), we performed an enrichment analysis on GO terms for biological processes based on a classic Fisher algorithm. Enriched GO terms with the lowest *p-value* were selected for representation. Dot size represent the number of genes belonging to the clusters associated with the GO term. (b) Average *z-score* values for each cluster. The coloured dotted line corresponds to the estimated date of dormancy release.

**Figure 5.**
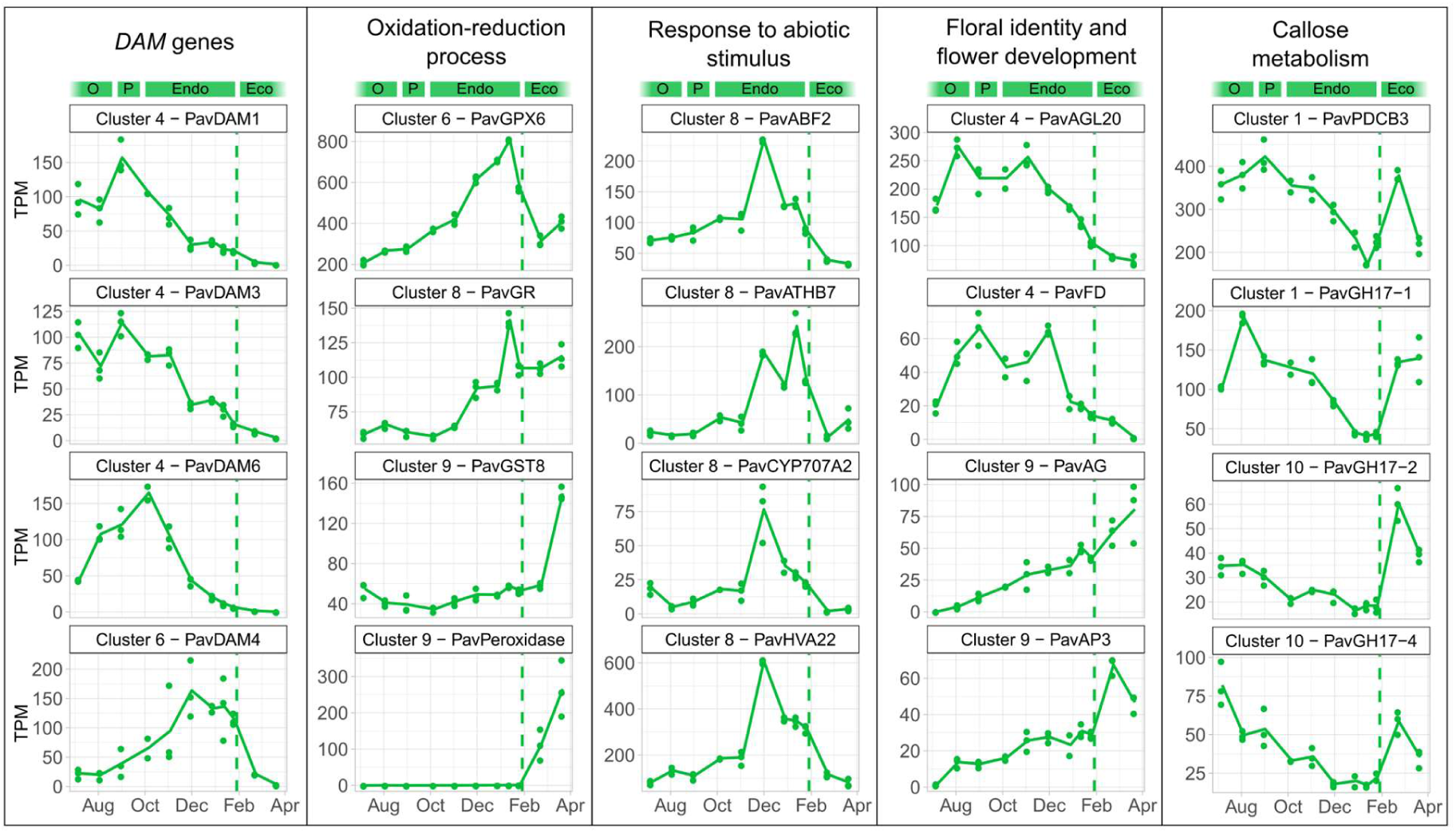
Expression patterns of key genes involved in sweet cherry bud dormancy. Expression patterns, expressed in transcripts per million reads (TPM) were analysed for the cultivar ‘Garnet’ from August to March, covering bud organogenesis (O), paradormancy (P), endodormancy (Endo), and ecodormancy (Eco). Dash lines represent the estimated date of dormancy breaking.

We further investigated whether dormancy-associated genes were specifically activated and repressed during the different bud stages. Among the six annotated *DAM* genes, four were differentially expressed in the dataset. *PavDAM1, PavDAM3* and *PavDAM6* were highly expressed during paradormancy and at the beginning of endodormancy (cluster 4, Fig. 5) whereas the expression peak for *PavDAM4* was observed at the end of endodormancy (cluster 6, Fig. 5). In addition, we found that genes coding for 1,3-β-glucanases from the Glycosyl hydrolase family 17 (*PavGH17*), as well as a *PLASMODESMATA CALLOSE-BINDING PROTEIN 3 (PavPDCB3)* gene were repressed during dormancy (clusters 1 and 10, Fig. 5).

### Specific transcription factor target genes are expressed during the main flower bud stages

To better understand the regulation of genes that are expressed at different flower bud stages, we investigated whether some transcription factors (TFs) targeted genes in specific clusters. Based on a list of predicted regulation between TFs and target genes that is available for peach in PlantTFDB [35], we identified the TFs with enriched targets in each cluster (Table 1). We further explored these target genes and their biological functions with a GO enrichment analysis (Tables S3, S4 in Additional file 2). Moreover, to have a complete overview of the TFs’ targets, we also identified enriched target promoter motifs in the different gene clusters (Table S5 in Additional file 2), using motifs we discovered with Find Individual Motif occurrences (FIMO) [36] and reference motifs obtained from PlantTFDB 4.0 [35]. Results show that different pathways are activated throughout bud development.

Among the genes expressed during the organogenesis and paradormancy phases (clusters 1, 2, 3 and 4), we observed an enrichment for motifs targeted by several MADS-box TFs such as AGAMOUS (AG), APETALA3 (AP3) and SEPALLATA3/AGAMOUS-like 9 (SEP3/AGL9), several of them potentially involved in flower organogenesis [37]. On the other hand, for the same clusters, results show an enrichment in MYB-related targets, WRKY and ethylene-responsive element (ERF) binding TFs (Table 1, Table S5 in Additional file 2). Several members of these TF families have been shown to participate in the response to abiotic factors. Similarly, we found in the cluster 4 target motifs enriched for DEHYDRATION RESPONSE ELEMENT-BINDING2 (PavDREB2C), potentially involved in the response to cold [38]. *PavMYB63* and *PavMYB93* transcription factors, expressed during organogenesis and paradormancy, likely activate genes involved in secondary metabolism (Table 1, Tables S3, S4 in Additional file 2).

During endodormancy, we found that *PavMYB14* and *PavMYB40* specifically target genes from cluster 10 that are involved in secondary metabolic processes and growth (Tables S3, S4 in Additional file 2). Expression profiles suggest that *PavMYB14* and *PavMYB40* repress expression of these target genes during endodormancy (Fig. S4 in Additional file 1). This is consistent with the functions of *Arabidopsis thaliana* MYB14 that negatively regulates the response to cold [39]. One of the highlighted TFs was *PavWRKY40*, which is activated before endodormancy and preferentially regulates genes associated with oxidative stress (Fig. S4 and Table S4 in Additional files 1 and 2).

Interestingly, we observed a global response to cold and stress during endodormancy since we identified an enrichment of targets for *PavCBF4*, and of genes with motifs for several ethylene-responsive element binding TFs such as *PavDREB2C* in the cluster 5. We also observed an enrichment in the same cluster for genes with motifs for *PavABI5* (Table S5 in Additional file 2). All these TFs are involved in the response to cold, in agreement with the fact that genes in the cluster 5 are expressed during endodormancy. Genes belonging to the clusters 6, 7 and 8 are highly expressed during deep dormancy and we found targets and target motifs for many TFs involved in the response to abiotic stresses. For example, we found motifs enriched in the cluster 7 for many TFs of the C2H2 family, which are involved in the response to a wide spectrum of stress conditions, such as extreme temperatures, salinity, drought or oxidative stress (Table S5; [40, 41]). Similarly, in the cluster 8, we also identified an enrichment in targets and motifs of many genes involved in the response to ABA and to abiotic stimulus, such as *PavABF2, PavAREB3, PavABI5, PavDREB2C* and *PavERF110* (Tables S3, S4 in Additional file 2) [38, 42]. Their targets include ABA-related genes *HIGHLY ABA-INDUCED PP2C GENE 1 (PavHAI1), PavCYP707A2* that is involved in ABA catabolism, *PavPYL8* a component of ABA receptor 3 and *LATE EMBRYOGENESIS ABUNDANT PROTEIN (PavLEA)*, involved in the response to desiccation [4].

We also observe during endodormancy an enrichment for targets of TFs involved in the response to light and temperature, such as *PavPIL5, PavSPT, PavRVE1* and *PavPIF4* (Table 1, [5, 43–45]), and *PavRVE8* that preferentially target genes involved in cellular transport like *LIPID TRANSFER PROTEIN1* (*PavLP1*, Table S3 in Additional file 2). Interestingly, we found that among the TFs with enriched targets in the clusters, only ten display changes in expression during flower bud development (Table 1, Fig. S4 in Additional file 1), including *PavABF2, PavABI5* and *PavRVE1*. Expression profiles for these three genes are very similar, and are also similar to their target genes, with a peak of expression around the estimated dormancy release date, indicating that these TFs are positively regulating their targets (see Fig. S4 in Additional file 1).

### Expression patterns highlight bud dormancy similarities and disparities between three cherry tree cultivars

Since temperature changes and progression through the flower bud stages are happening synchronously, it is challenging to discriminate transcriptional changes that are mainly associated with one or the other. In this context, we also analysed the transcriptome of two other sweet cherry cultivars: ‘Cristobalina’, characterized by very early flowering dates, and ‘Regina’, with a late flowering time. The span between flowering periods for the three cultivars is also found in the transition between endodormancy and ecodormancy since ten weeks separated the estimated dates of dormancy release between the cultivars: 9th December 2015 for ‘Cristobalina’, 29th January 2016 for ‘Garnet’ and 26th February 2016 for ‘Regina’ (Fig. 1a). The transition from organogenesis to paradormancy is not well documented and many studies suggest that endodormancy onset is under the strict control of environment in *Prunus* species [3]. Therefore, we considered that these two transitions occurred at the same time in all three cultivars. However, the two months and half difference in the date of transition from endodormancy to ecodormancy between the cultivars allow us to look for transcriptional changes associated with this transition independently of environmental conditions. To do so, we generated a total of 50 transcriptomes from buds harvested at ten dates for the cultivar ‘Cristobalina’, and eleven dates for the cultivar ‘Regina’, spanning all developmental stages from bud organogenesis to flowering. We then compared the expression patterns between the three contrasted cultivars throughout flower bud stages for the genes we identified as differentially expressed in the cultivar ‘Garnet’ (Fig. 1b).

When projected into a PCA 2-components plane, all samples harvested from buds at the same stage cluster together, whatever the cultivar (Fig. 6 and Fig. S5), suggesting that the stage of the bud has more impact on the transcriptional state than time or external conditions. Interestingly, the 100 genes that contributed the most to the PCA dimensions 1 and 2 were very specifically associated with each dimension (Fig. S6, Table S6). We further investigated which clusters were over-represented in these genes (see Fig. S6b in Additional file 1) and we found that genes belonging to the clusters 6 and 8, associated with endodormancy, were particularly represented in the best contributors to the dimension 1. In particular, we identified genes involved in oxidation-reduction processes like *PavGPX6*, and stress-induced genes such as *PavLEA14*, together with genes potentially involved in leaf and flower development, including *GROWTH-REGULATING FACTOR7 (PavGRF7)* and *PavSEP1* (Table S6). In contrast, genes that best contributed to the dimension 2 strictly belonged to clusters 9 and 10, therefore characterized by high expression during ecodormancy (Fig. S6 in Additional file 1). These results suggest that bud stages can mostly be separated by two criteria: dormancy depth before dormancy release, defined by genes highly expressed during endodormancy, and the dichotomy defined by the status before/after dormancy release.

**Figure 6.**
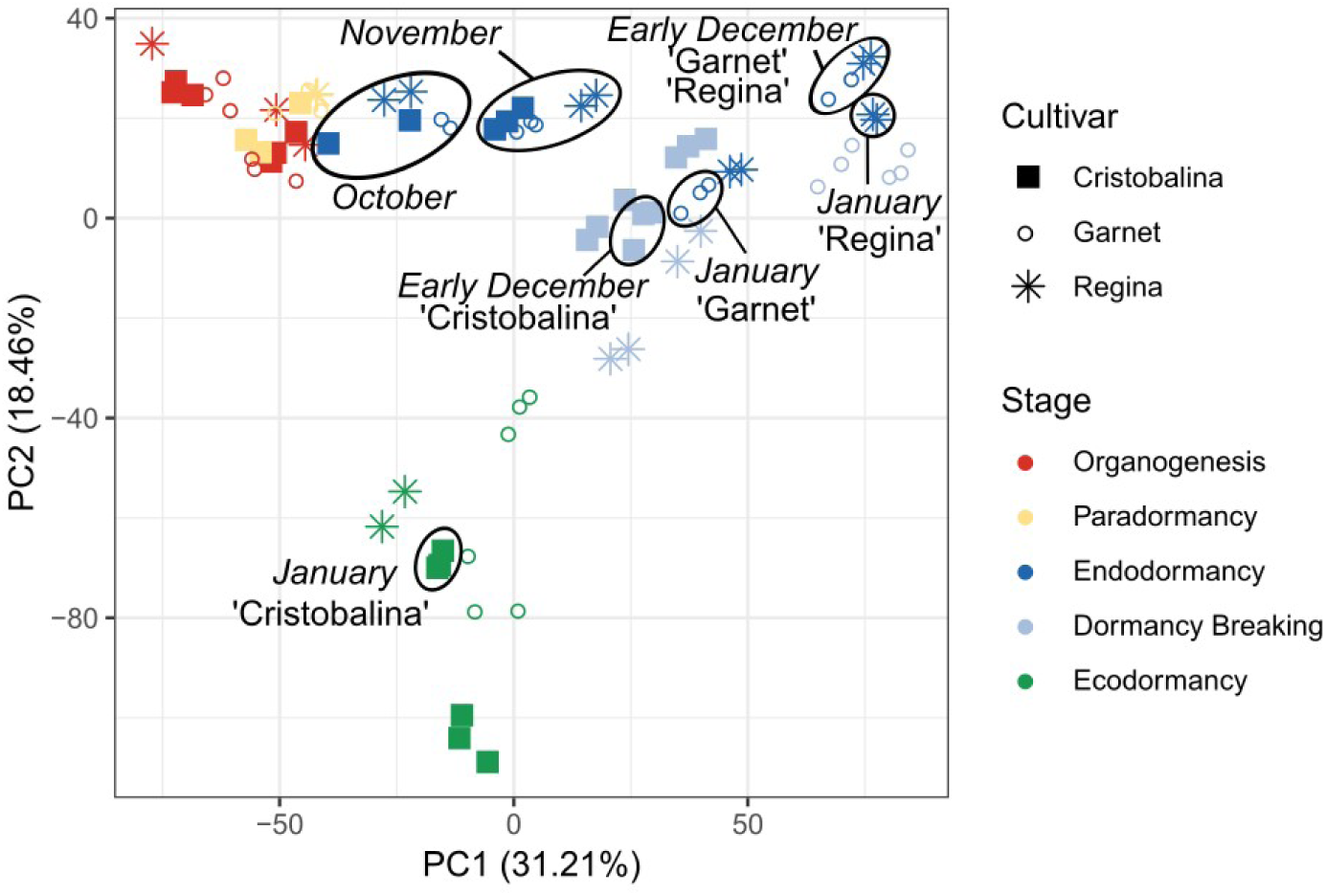
Separation of samples by dormancy stage and cultivar using differentially expressed genes. The principal component analysis was conducted on the TPM (transcripts per millions reads) values for the differentially expressed genes in the flower buds of the cultivars ‘Cristobalina’ (filled squares), ‘Garnet’ (empty circles) and ‘Regina’ (stars). Each point corresponds to one sampling time in a single tree.

To go further, we compared transcriptional profiles throughout the time course in all cultivars. For this we analysed the expression profiles in each cultivar for the clusters previously identified for the cultivar ‘Garnet’ (Fig. 7, see also Fig. S7). In general, averaged expression profiles for all clusters are very similar in all three varieties, with the peak of expression happening at a similar period of the year. However, we can distinguish two main phases according to similarities or disparities between cultivars. First, averaged expression profiles are almost similar in all cultivars between July and November. This is especially the case for clusters 1, 4, 7, 8 and 9.

**Figure 7.**
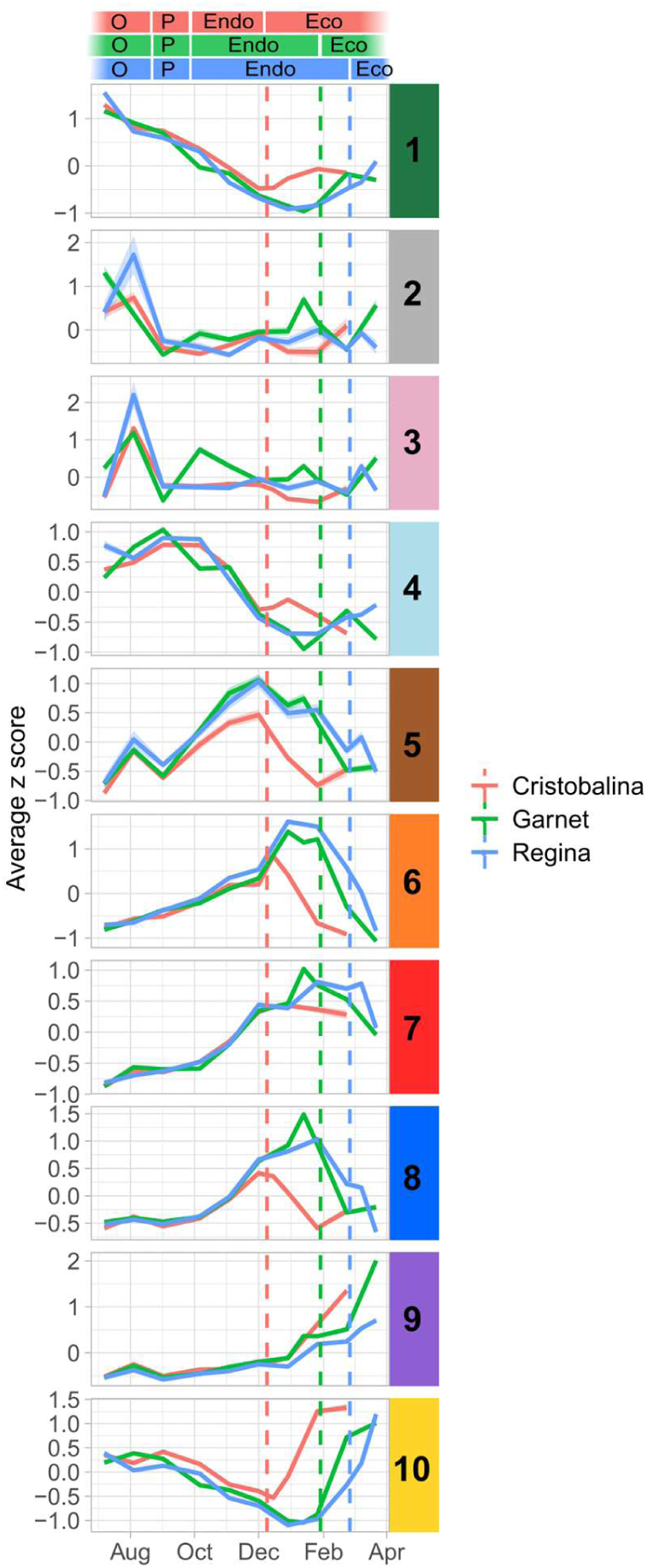
Expression patterns in the ten clusters for the three cultivars. Expression patterns were analysed from August to March, covering bud organogenesis (O), paradormancy (P), endodormancy (Endo), and ecodormancy (Eco). Dash lines represent the estimated date of dormancy breaking, in red for ‘Cristobalina’, green for ‘Garnet’ and blue for ‘Regina’. Average *z-score* patterns (line) and standard deviation (ribbon), calculated using the TPM values from the RNA-seq analysis, for the genes belonging to the ten clusters.

On the other hand, we can observe a temporal shift in the peak of expression between varieties from December onward for genes in clusters 1, 5, 6, 8 and 10. Indeed, in these clusters, the peak or drop in expression happens earlier in ‘Cristobalina’, and slightly later in ‘Regina’ compared to ‘Garnet’ (Fig. 7), in correlation with their dormancy release dates. These results seem to confirm that the organogenesis and paradormancy phases occur concomitantly in the three cultivars while temporal shifts between cultivars are observed after endodormancy onset. Therefore, similarly to the PCA results (Fig. 6), the expression profile of these genes is more associated with the flower bud stage than with external environmental conditions.

### Flower bud stage can be predicted using a small set of marker genes

We have shown that flower buds in organogenesis, paradormancy, endodormancy and ecodormancy are characterised by specific transcriptional states. In theory, we could therefore use transcriptional data to infer the flower bud stage. For this, we selected a minimum number of seven marker genes, one gene for each of the clusters 1, 4, 5, 7, 8, 9 and 10 (identified in Fig 3), for which expression presented the best correlation with the average expression profiles of their cluster (Fig. 8). We aimed to select the minimum number of marker genes that are sufficient to infer the flower bud stage, therefore excluding the clusters 2, 3 and 6 as they either had very small number of genes, or had expression profiles very similar to another cluster.

**Figure 8.**
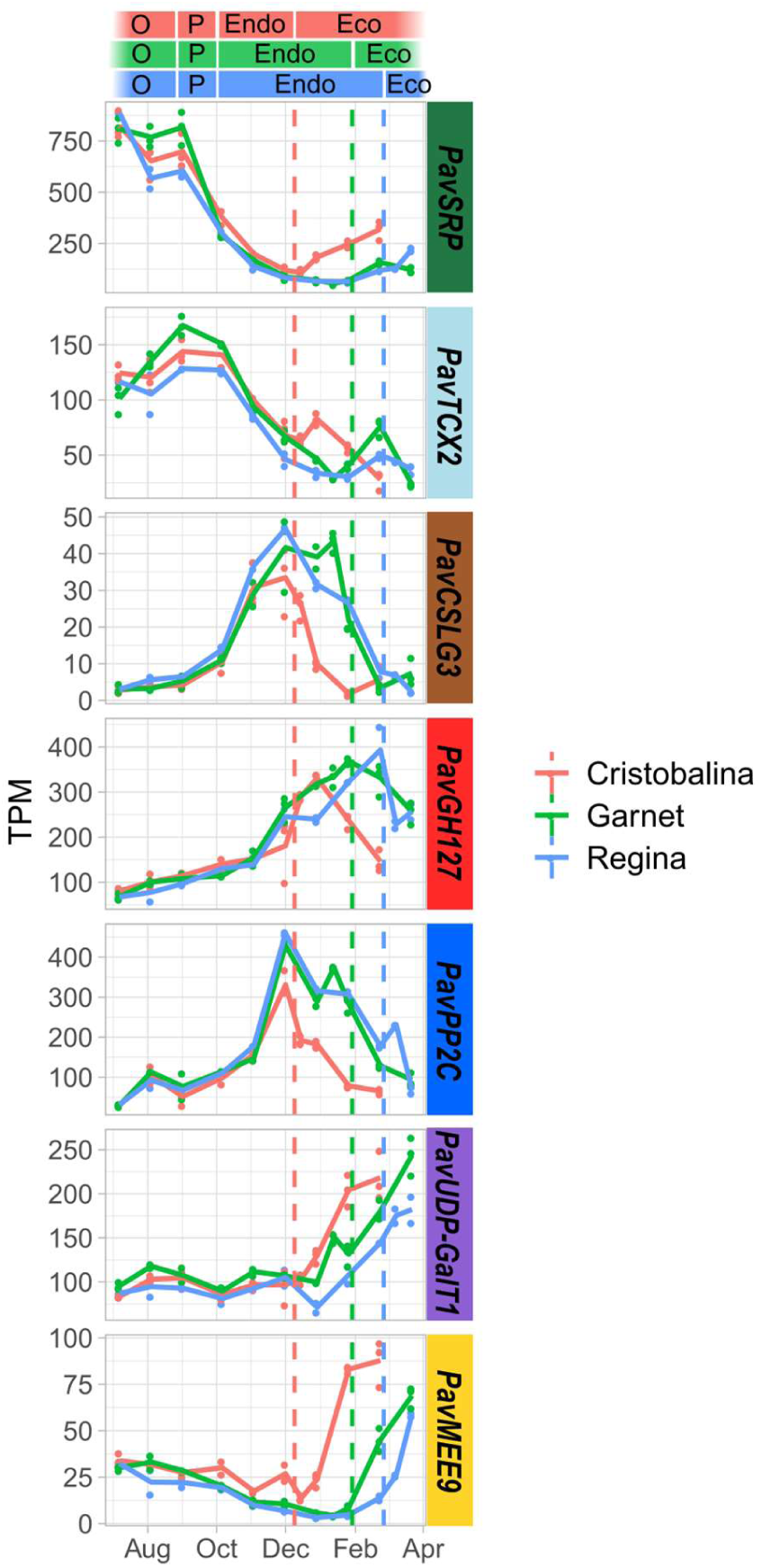
Expression patterns for the seven marker genes in the three cultivars. Expression patterns were analysed from August to March, covering bud organogenesis (O), paradormancy (P), endodormancy (Endo), and ecodormancy (Eco). Dash lines represent the estimated date of dormancy breaking, in red for ‘Cristobalina’, green for ‘Garnet’ and blue for ‘Regina’. TPM were obtained from the RNA-seq analysis for the seven marker genes from clusters 1, 4, 5, 7, 8, 9 and 10. Lines represent the average TPM, dots are the actual values from the biological replicates. *SRP: STRESS RESPONSIVE PROTEIN; TCX2: TESMIN/TSO1-like CXC 2; CSLG3: Cellulose Synthase like G3; GH127: Glycosyl Hydrolase 127; PP2C: Phosphatase 2C; UDP-GalT1: UDP-Galactose transporter 1; MEE9: maternal effect embryo arrest 9*.

Expression for these marker genes not only recapitulates the average profile of the cluster they originate from, but also temporal shifts in the profiles between the three cultivars (Fig. 8). In order to define if these genes encompass as much information as the full transcriptome, or all DEGs, we performed a PCA of all samples harvested for all three cultivars using expression levels of these seven markers (Fig. S9). The clustering of samples along the two main axes of the PCA using these seven markers is very similar, if not almost identical, to the PCA results obtained using expression for all DEGs (Fig. 6). This indicates that the transcriptomic data can be reduced to only seven genes and still provides accurate information about the flower bud stages.

To test if these seven markers can be used to define the flower bud stage, we used a multinomial logistic regression modelling approach to predict the flower bud stage in our dataset based on the expression levels for these seven genes in the three cultivars ‘Garnet’, ‘Regina’ and ‘Cristobalina’ (Fig. 9). For this, we trained and tested the model, on randomly picked sets, to predict the five bud stage categories, and obtained a very high model accuracy (100%; Fig. S9). These results indicate that the bud stage can be accurately predicted based on expression data by just using seven genes. In order to go further and test the model in an independent experiment, we analysed the expression for the seven marker genes by RT-qPCR on buds sampled from another sweet cherry tree cultivar ‘Fertard’ for two consecutive years (Fig. 9a, b). Based on these RT-qPCR data, we predicted the flower bud developmental stage using the parameters of the model obtained from the training set on the three cultivars ‘Garnet’, ‘Regina’ and ‘Cristobalina’. We achieved a high accuracy of 71% for our model when tested on RT-qPCR data to predict the flower bud stage for the ‘Fertard’ cultivar (Fig. 9c and Fig. S9c). In particular, the chronology of bud stages was very well predicted. This result indicates that these seven genes can be used as a diagnostic tool in order to infer the flower bud stage in sweet cherry trees.

**Figure 9.**
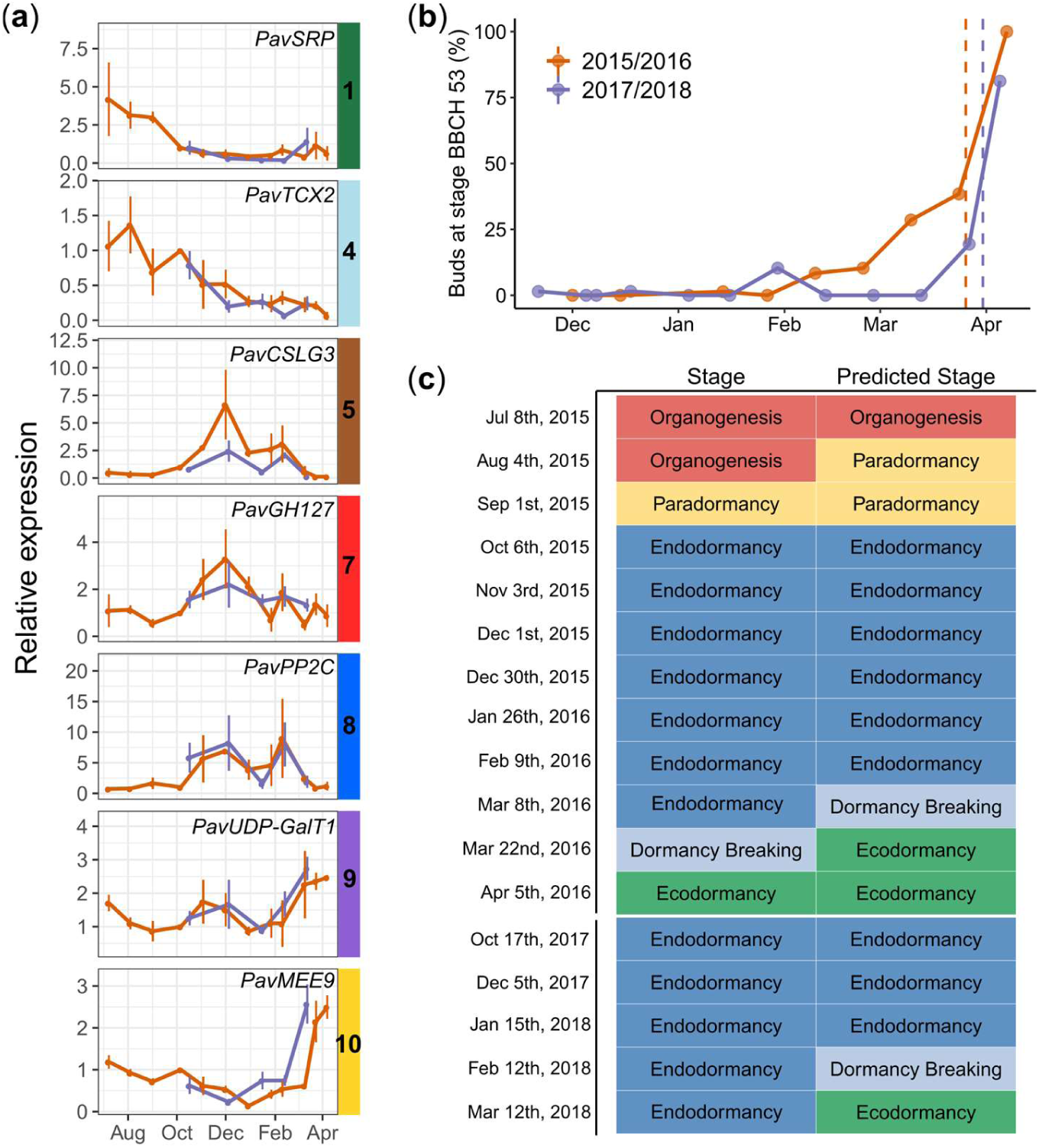
Expression for the seven marker genes allows accurate prediction of the bud dormancy stages in the late flowering cultivar ‘Fertard’ during two bud dormancy cycles Relative expressions were obtained by qRT-PCR and normalized by the expression of two reference constitutively expressed genes *PavRPII* and *PavEF1*. (b) Evaluation of the dormancy status in ‘Fertard’ flower buds during the two seasons using the percentage of open flower buds (BBCH stage 53). (c) Predicted vs experimentally estimated bud stages. *SRP: STRESS RESPONSIVE PROTEIN; TCX2: TESMIN/TSO1-like CXC 2; CSLG3: Cellulose Synthase like G3; GH127: Glycosyl Hydrolase 127; PP2C: Phosphatase 2C; UDP-GalT1: UDP-Galactose transporter 1; MEE9: maternal effect embryo arrest 9*.

## DISCUSSION

In this work, we have characterised transcriptional changes at a genome-wide scale happening throughout cherry tree flower bud dormancy, from organogenesis to the end of dormancy. To do this, we have analysed expression in flower buds at 11 dates from July 2015 to March 2016 for three cultivars displaying different dates of dormancy release, generating 82 transcriptomes in total. This resource, with a fine time resolution, reveals key aspects of the regulation of cherry tree flower buds during dormancy (Fig. 10). We have shown that buds in organogenesis, paradormancy, endodormancy and ecodormancy are characterised by distinct transcriptional states (Fig. 2, 3) and we highlighted the different pathways activated during the main cherry tree flower bud dormancy stages (Fig. 4 and Table 1). Finally, we found that just seven genes are enough to accurately predict the main cherry tree flower bud dormancy stages (Fig. 9).

**Figure 10.**
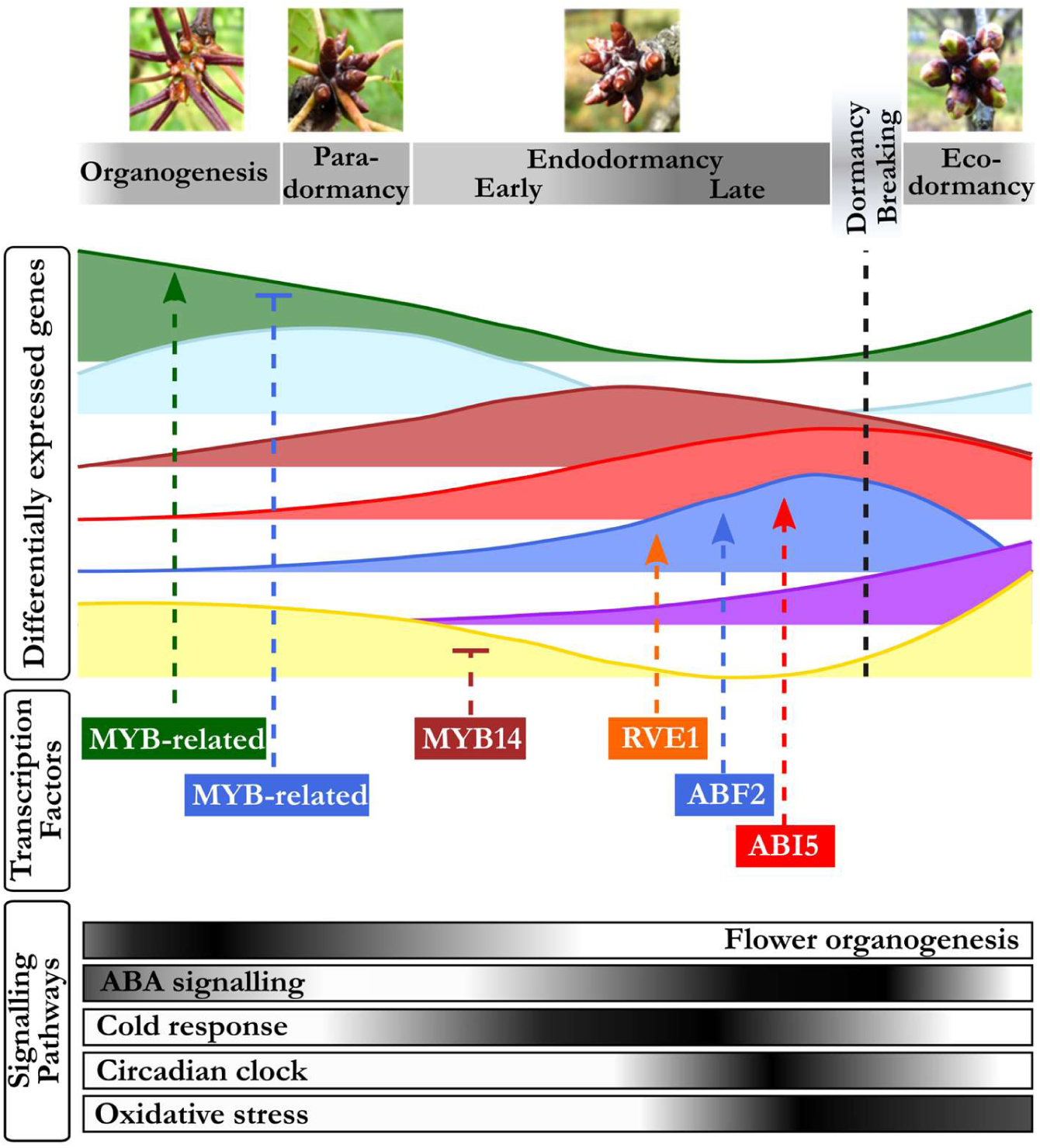
From bud formation to flowering: transcriptomic regulation of flower bud dormancy. Our results highlighted seven main expression patterns corresponding to the main dormancy stages. During organogenesis and paradormancy (July to September), signalling pathways associated with flower organogenesis and ABA signalling are upregulated. Distinct groups of genes are activated during different phases of endodormancy, including targets of transcription factors involved in ABA signalling, cold response and circadian clock. ABA: abscisic acid.

### *DAMs*, floral identity and organogenesis genes characterize the pre-dormancy stages

To our knowledge, this is the first report on the transcriptional regulation of early stages of flower bud development in temperate fruit trees. Information on dormancy onset and pre-dormancy bud stages are scarce and we arbitrarily delimited the organogenesis and paradormancy in July/August and September, respectively. However, based on transcriptional data, we could detect substantial discrepancies suggesting that the definition of the bud stages can be improved. Indeed, we observe that samples harvested from buds during phases that we defined as organogenesis and paradormancy cluster together in the PCA, but away from samples harvested during endodormancy. Moreover, most of the genes highly expressed during paradormancy are also highly expressed during organogenesis. This is further supported by the fact that paradormancy is a flower bud stage predicted with less accuracy based on expression level of the seven marker genes. In details, paradormancy is defined as a stage of growth inhibition originating from surrounding organs [7] therefore it is strongly dependent on the position of the buds within the tree and the branch. Our results suggest that defining paradormancy for multiple cherry flower buds based on transcriptomic data is difficult and even raise the question of whether paradormancy can be considered as a specific flower bud stage. Alternatively, we propose that the pre-dormancy period should rather be defined as a continuum between organogenesis, growth and/or growth cessation phases. Further physiological observations, including flower primordia developmental context [46], could provide crucial information to precisely link the transcriptomic environment to these bud stages. Nonetheless, we found very few, if not at all, differences between the three cultivars for the expression patterns during organogenesis and paradormancy, supporting the hypothesis that pre-dormancy processes are not associated with the different timing in dormancy release and flowering that we observe between these cultivars.

Our results showed that specific pathways were specifically activated before dormancy onset. The key role of ABA in the control of bud set and dormancy onset has been known for decades and we found that the ABA-related transcription factor *PavWRKY40* is expressed as early as during organogenesis. Several studies have highlighted a role of *PavWRKY40* homolog in Arabidopsis in ABA signalling, in relation with light transduction [47, 48] and biotic stresses [49]. These results suggest that there might be an early response to ABA in flower buds. Furthermore, we uncovered the upregulation of several pathways linked to organogenesis during the summer months, including *PavMYB63* and *PavMYB93*, expressed during early organogenesis, with potential roles in the secondary wall formation [50] and root development [51]. Interestingly, *TESMIN/TSO1-like CXC 2 (PavTCX2)*, defined here as a marker gene for organogenesis and paradormancy, is the homolog of an Arabidopsis TF potentially involved in stem cell division [52]. We found that targets for *PavTCX2* may be over-represented in genes up-regulated during endodormancy, thus suggesting that *PavTCX2* acts on bud development by repressing dormancy-associated genes. In accordance with the documented timing of floral initiation and development in sweet cherry [53], several genes involved in floral identity and flower development, including *PavAGL20, PavFD*, as well as targets of *PavSEP3, PavAP3* and *PavAG*, were markedly upregulated during the early stages of flower bud development. Many studies conducted on fruit trees support the key role of *DAM* genes in the control of dormancy establishment and maintenance [18] and we found expression patterns very similar to the peach *DAM* genes with *PavDAM1* and *PavDAM3*, as well as *PavDAM6*, expressed mostly during summer [54]. The expression of these three genes was at the highest before endodormancy and seems to be inhibited by cold exposure from October onward, similarly to previous results obtained in sweet cherry [55], peach [56], Japanese apricot [57] and apple [58]. These results further suggest a major role for *PavDAM1, PavDAM3* and *PavDAM6* in dormancy establishment, bud onset and growth cessation in sweet cherry.

### Integration of environmental and internal signals through a complex array of signaling pathways during endodormancy

Previous studies have proved the key role of a complex array of signaling pathways in the regulation of endodormancy onset and maintenance that subsequently lead to dormancy breaking, including genes involved in cold response, phytohormone-associated pathways and oxidation-reduction processes. Genes associated with the response to cold, notably, have been shown to be up-regulated during endodormancy such as dehydrins and *DREB* genes identified in oak, pear and leafy spurge [24, 27, 59]. We observe an enrichment for GO involved in the response to abiotic and biotic responses, as well as an enrichment for targets of many TFs involved in the response to environmental factors. In particular, our results suggest that *PavMYB14*, which has a peak of expression in November just before the cold period starts, is repressing genes that are subsequently expressed during ecodormancy. This is in agreement with the fact that *AtMYB14*, the *PavMYB14* homolog in Arabidopsis thaliana, is involved in cold stress response regulation [39]. Although these results were not confirmed in *Populus* [60], two MYB DOMAIN PROTEIN genes (*MYB4* and *MYB14*) were also up-regulated during the induction of dormancy in grapevine [61]. Similarly, we identified an enrichment in genes highly expressed during endodormancy with target motifs of a transcription factor belonging to the CBF/DREB family. These TFs have previously been implicated in cold acclimation and endodormancy in several perennial species [59, 62]. These results are in agreement with the previous observation showing that genes responding to cold are differentially expressed during dormancy in other tree species [24]. Cold acclimation is the ability of plants to adapt to and withstand freezing temperatures and is triggered by decreasing temperatures and photoperiod. Therefore mechanisms associated with cold acclimation are usually observed concomitantly to the early stages of endodormancy. The stability of membranes and a strict control of cellular homeostasis are crucial in the bud survival under cold stress and we observe that genes associated with cell wall organization and nutrient transporters are up-regulated at the beginning of endodormancy, including the *CELLULOSE SYNTHASE-LIKE G3 (PavCSLG3)* marker gene.

Similarly to seed dormancy processes, hormonal signals act in a complex way to balance dormancy maintenance and growth resumption. In particular, ABA levels have been shown to increase in response to environmental signals such as low temperatures and/or shortening photoperiod, and trigger dormancy induction [63–65] Several studies have also shown that a subsequent drop in ABA concentration is associated with dormancy release [64, 66]. These results are supported by previous reports where genes involved in ABA signaling are differentially expressed during dormancy in various tree species (for e.g., see [19, 20, 22, 24, 67]). We find ABA-related pathways to be central in our transcriptomic analysis of sweet cherry bud dormancy, with the enrichment of GO terms related to ABA found in the genes highly expressed during endodormancy. These genes, including ABA-degradation gene *PavCYP707A2*, ABA-response factor *PavABF2*, and the Protein phosphatase 2C (PavPP2C) marker gene, are then inhibited after dormancy release in the three cultivars. Accordingly, we identified a key role for ABA-associated genes *PavABI5* and *PavABF2* in the regulation of dormancy progression in our dataset. These two transcription factors are mainly expressed around the time of dormancy release, like their target, and their homologs in Arabidopsis are involved in key ABA processes, especially during seed dormancy [68]. These results are consistent with records that *PmABF2* is highly expressed during endodormancy in Japanese apricot [22]. These results suggest ABA-related mechanisms similar in sweet cherry to those previously observed in other trees control bud dormancy onset and release. One of the hypotheses supports an activation of ABA-induced dormancy by *DAM* genes [64, 69] and we observed that *PavDAM4* expression pattern is very similar to ABA-related genes. We can therefore hypothesize that *PavDAM4* has a key role in dormancy onset and maintenance, potentially by regulating ABA metabolism. On the other side of the pathway, ground-breaking works have revealed that ABA signaling is crucial in triggering dormancy onset by inducing plasmodesmata closure, potentially through callose deposit [65, 70]. Accordingly, we found that *PavGH17* genes involved in callose degradation are highly activated before and after endodormancy while their expression is inhibited during endodormancy, thus suggesting that callose deposit is activated during endodormancy in sweet cherry flower buds.

In plants, response to environmental and developmental stimuli usually involves pathways associated with circadian clock regulation. This is also true for bud dormancy where the interplay between environmental and internal signals necessitates circadian clock genes for an optimal response [4, 71–74]. Indeed, transcriptomic analyses conducted in poplar showed that among the genes up-regulated during endodormancy, were genes with the *EVENING ELEMENT (EE)* motifs, that are important regulators of circadian clock and cold-responsive genes, and components of the circadian clock, including *LATE-ELONGATE HYPOCOTYL (LHY) and ZEITLUPE (ZTL)* [60, 67]. We identified an enrichment of targets for *PavRVE8* and *PavRVE1* among the genes expressed around the time of dormancy release. Homologs of RVE1 are also up-regulated during dormancy in leafy spurge [43] and apple [75]. These TFs are homologs of Arabidopsis MYB transcription factors involved in the circadian clock. In particular, *AtRVE1* seems to integrate several signalling pathways including cold acclimation and auxin [76–78] while *AtRVE8* is involved in the regulation of circadian clock by modulating the pattern of H3 acetylation [79]. Our findings that genes involved in the circadian clock are expressed and potentially regulate genes at the time of dormancy release strongly support the hypothesis that environmental cues might be integrated with internal factors to control dormancy and growth in sweet cherry flower buds.

Consistently with observations that elevated levels of the reactive species of oxygen H_2_O_2_ are strongly associated with dormancy release [80], oxidative stress is considered as one of the important processes involved in the transition between endodormancy and ecodormancy [30, 81, 82]. In line with these findings, we identified genes involved in oxidation-reduction processes that are up-regulated just before endodormancy release including *PavGPX6* and *PavGR*, that are involved in the detoxification systems. In their model for the control of dormancy, Ophir and colleagues [82] hypothesize that respiratory stress, ethylene and ABA pathways interact to control dormancy release and growth resumption. Our results concur with this hypothesis to some extend albeit the key role of *DAM* genes should be further explored. Co-regulation analyses will be needed to investigate whether oxidative stress signalling is involved upstream to trigger dormancy release or downstream as a consequence of cellular activity following dormancy release in sweet cherry buds, leading to a better understanding of how other pathways interact or are directly controlled by oxidative cues.

### Global cell activity characterizes the ecodormancy stage in sweet cherry flower buds

Following the release of endodormancy, buds enter the ecodormancy stage, which is a state of inhibited growth controlled by external signals that can therefore be reversed by exposure to growth-promoting signals [7]. This transition towards the ability to grow is thought to be associated with the prolonged downregulation of *DAM* genes (see [18] for review), regulated by epigenetic mechanisms such as histone modifications [62, 83–85] and DNA methylation [55], in a similar way to *FLC* repression during vernalization in Arabidopsis. We observe that the expression of all *PavDAM* genes is inhibited before dormancy release, thus supporting the hypothesis that *DAM* genes may be involved in dormancy maintenance. In particular, the transition to ecodormancy coincides with a marked decrease in *PavDAM4* expression, which suggest that the regulation of its expression is crucial in the progression of dormancy towards growth resumption. However, other MADS-box transcription factors were found to be up-regulated during ecodormancy, including *PavAG* and *PavAP3*, similarly to previous results obtained in Chinese cherry (*Prunus pseudocerasus*) [28]. We also found that the marker gene *PavMEE9*, expressed during ecodormancy, is orthologous to the Arabidopsis gene *MATERNAL EFFECT EMBRYO ARREST 9 (MEE9)*, required for female gametophyte development [86], which could suggest active cell differentiation during the ecodormancy stage.

As mentioned before, in-depth studies conducted on poplar have led to the discovery that the regulation of the movements through the plasma membrane plays a key role not only in dormancy onset but also in dormancy release [87]. This is also true for long-distance transport with the observation that in peach, for example, active sucrose import is renewed during ecodormancy [88]. In sweet cherry, our results are consistent with these processes since we show that GO terms associated with transmembrane transporter activity are enriched for genes highly expressed during ecodormancy. Transmembrane transport capacity belongs to a wide range of membrane structures modifications tightly regulated during dormancy. For example, lipid content, linoleic and linolenic acids composition and unsaturation degree of fatty acids in the membrane are modified throughout dormancy progression [30] and these changes in the membrane structure may be associated with modifications in the cytoskeleton [87]. Consistently, we find that genes involved in microtubule-based processes and cell wall organization are up-regulated during ecodormancy in sweet cherry flower buds. For example, the marker gene *PavUDP-GalT1*, orthologous to a putative UDP-galactose transmembrane transporter, is highly express after dormancy release in all three cultivars.

Overall, all processes triggered during ecodormancy are associated with cell activity. The trends observed here suggest that after endodormancy release, transmembrane and long distance transports are reactivated, thus allowing an active uptake of sugars, leading to increased oxidation-reduction processes and cell proliferation and differentiation.

### Development of a diagnostic tool to define the flower bud dormancy stage using seven genes

We find that sweet cherry flower bud stage can be accurately predicted with the expression of just seven genes. It indicates that combining expression profiles of just seven genes is enough to recapitulate all transcriptional states in our study. This is in agreement with previous work showing that transcriptomic states can be accurately predicted using a relatively low number of markers [89]. Marker genes were not selected on the basis of their function and indeed, two genes are orthologous to Arabidopsis proteins of unknown function: *PavSRP* (Stress responsive A/B Barrel Domain-containing protein) and *PavGH127* (putative glycosyl hydrolase). However, as reported above, some of the selected marker genes are involved in the main pathways regulating dormancy progression, including cell wall organization during the early phase of endodormancy (*PavCSLG3*), ABA (*PavPP2C*), transmembrane transport (*PavUDP-GalT1*) and flower primordia development (*PavMEE9*).

Interestingly, when there are discrepancies between the predicted bud stages and the ones defined by physiological observations, the model always predicts that stages happen earlier than the actual observations. For example, the model predicts that dormancy breaking occurs instead of endodormancy, or ecodormancy instead of dormancy breaking. This could suggest that transcriptional changes happen before we can observe physiological changes. This is indeed consistent with the indirect phenotyping method currently used, based on the observation of the response to growth-inducible conditions after ten days. Using these seven genes to predict the flower bud stage would thus potentially allow to identify these important transitions when they actually happen.

We show that the expression level of these seven genes can be used to predict the flower bud stage in other conditions and genotypes by performing RT-qPCR. Also this independent experiment has been done on two consecutive years and shows that RT-qPCR for these seven marker genes as well as two control genes are enough to predict the flower bud stage in cherry trees. It shows that performing a full transcriptomic analysis is not necessary if the only aim is to define the dormancy stage of flower buds.

## CONCLUSIONS

In this work, we have characterized transcriptional changes throughout all stages of sweet cherry flower bud development and dormancy. To our knowledge, no analysis had previously been conducted on this range of dates in temperate trees. Pathways involved at different stages of bud dormancy have been investigated in other species and we confirmed that genes associated with the response to cold, ABA and development processes were also identified during sweet cherry flower bud dormancy. We took advantage of the extended timeframe and we highlighted genes and pathways associated with specific phases of dormancy, including early endodormancy, deep endodormancy and dormancy release. For that reason, our results suggest that commonly used definitions of bud dormancy are too restrictive and transcriptomic states might be useful to redefine the dormancy paradigm, not only for sweet cherry but also for other species that undergo overwintering. We advocate for large transcriptomic studies that take advantage of the wide range of genotypes available in forest and fruit trees, aiming at the mechanistic characterization of dormancy stages. Furthermore, we then went a step beyond the global transcriptomic analysis and we developed a model based on the transcriptional profiles of just seven genes to accurately predict the main dormancy stages. This offers an alternative approach to methods currently used such as assessing the date of dormancy release by using forcing conditions. In addition, this result sets the stage for the development of a fast and cost effective diagnostic tool to molecularly define the dormancy stages in cherry trees. This approach, from transcriptomic data to modelling, could be tested and transferred to other fruit tree species and such diagnostic tool would be very valuable for researchers working on fruit trees as well as for plant growers, notably to define the best time for the application of dormancy breaking agents, whose efficiency highly depends on the state of dormancy progression.

## METHODS

### Plant material

Branches and flower buds were collected from four different sweet cherry cultivars with contrasted flowering dates: ‘Cristobalina’, ‘Garnet’, ‘Regina’ and ‘Fertard’, which display extra-early, early, late and very late flowering dates, respectively. ‘Cristobalina’, ‘Garnet’, ‘Regina’ trees were grown in an orchard located at the Fruit Experimental Unit of INRA in Bourran (South West of France, 44° 19′ 56′′ N, 0° 24′ 47′′ E), under the same agricultural practices. ‘Fertard’ trees were grown in an orchard at the Fruit Experimental Unit of INRA in Toulenne, near Bordeaux (48° 51′ 46′′ N, 2° 17′ 15′′ E). During the first sampling season (2015/2016), ten or eleven dates spanning the entire period from flower bud organogenesis (July 2015) to bud break (March 2016) were chosen for RNA sequencing (Fig. 1a and Additional file 11), while bud tissues from ‘Fertard’ were sampled in 2015/2016 (12 dates) and 2017/2018 (7 dates) for validation by RT-qPCR (Additional file 11). For each date, flower buds were sampled from different trees, each tree corresponding to a biological replicate. Upon harvesting, buds were flash frozen in liquid nitrogen and stored at −80°C prior to performing RNA-seq.

### Measurements of bud break and estimation of the dormancy release date

For the two sampling seasons, 2015/2016 and 2017/2018, three branches bearing floral buds were randomly chosen fortnightly from ‘Cristobalina’, ‘Garnet’, ‘Regina’ and ‘Fertard’ trees, between November and flowering time (March-April). Branches were incubated in water pots placed under forcing conditions in a growth chamber (25°C, 16h light/ 8h dark, 60-70% humidity). The water was replaced every 3-4 days. After ten days under forcing conditions, the total number of flower buds that reached the BBCH stage 53 [46, 90] was recorded. The date of dormancy release was estimated as the date when the percentage of buds at BBCH stage 53 was above 50% after ten days under forcing conditions (Fig. 1a).

### RNA extraction and library preparation

Total RNA was extracted from 50-60 mg of frozen and pulverised flower buds using RNeasy Plant Mini kit (Qiagen) with minor modification: 1.5% PVP-40 was added in the extraction buffer RLT. RNA quality was evaluated using Tapestation 4200 (Agilent Genomics). Library preparation was performed on 1 μg of high quality RNA (RNA integrity number equivalent superior or equivalent to 8.5) using the TruSeq Stranded mRNA Library Prep Kit High Throughput (Illumina cat. no. RS-122-2103) for ‘Cristobalina’, ‘Garnet’ and ‘Regina’ cultivars. DNA quality from libraries was evaluated using Tapestation 4200. The libraries were sequenced on a NextSeq500 (Illumina), at the Sainsbury Laboratory Cambridge University (SLCU), using paired-end sequencing of 75 bp in length.

### Mapping and differential expression analysis

The raw reads obtained from the sequencing were analysed using several publicly available software and in-house scripts. The quality of reads was assessed using FastQC (www.bioinformatics.babraham.ac.uk/projects/fastqc/) and possible adaptor contaminations were removed using Trimmomatic [91]. Trimmed reads were mapped to the peach (*Prunus persica* (L) Batsch) reference genome v.2 [92] (genome sequence and information can be found at the following address: https://phytozome.jgi.doe.gov/pz/portal.html#!info?alias=Org_Ppersica) using Tophat [93].

Possible optical duplicates were removed using Picard tools (https://github.com/broadinstitute/picard). The total number of mapped reads of each samples are given in Additional file 15. For each gene, raw read counts and TPM (Transcripts Per Million) numbers were calculated [94].

We performed a differential expression analysis on data obtained from the ‘Garnet’ samples. First, data were filtered by removing lowly expressed genes (average read count < 3), genes not expressed in most samples (read counts = 0 in more than 75% of the samples); and genes presenting little change in expression between samples (coefficient of variation < 0.3). Then, differentially expressed genes (DEGs) between bud stages (organogenesis – 6 biological replicates, paradormancy – 3 biological replicates, endodormancy – 10 biological replicates, dormancy breaking – 6 biological replicates, ecodormancy – 6 biological replicates, see Additional file 11) were assessed using DEseq2 R Bioconductor package [95], in the statistical software R (R Core Team 2018), on filtered data. Genes with an adjusted *p-value* (padj) < 0.05, using the Benjamini-Hochberg multiple testing correction method, were assigned as DEGs (Additional file 12). To enable researchers to access this resource, we have created a graphical web interface to allow easy visualisation of transcriptional profiles throughout flower bud dormancy in the three cultivars for genes of interest (bwenden.shinyapps.io/DorPatterns/).

### Principal component analyses and hierarchical clustering

Distances between the DEGs expression patterns over the time course were calculated based on Pearson’s correlation on ‘Garnet’ TPM values. We applied a hierarchical clustering analysis on the distance matrix to define ten clusters (Additional file 12). For expression patterns representation, we normalized the data using *z-score* for each gene:

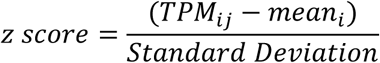

where TPM_ij_ is the TPM value of the gene *i* in the sample *j, mean_i_* and *standard deviationi* are the *mean* and *standard deviation* of the TPM values for the gene *i* over all samples.

Principal component analyses (PCA) were performed on TPM values from different datasets using the *prcomp* function from R.

For each cluster, using data for ‘Garnet’, ‘Regina’ and ‘Cristobalina’, mean expression pattern was calculated as the mean *z-score* value for all genes belonging to the cluster. We then calculated the Pearson’s correlation between the *z-score* values for each gene and the mean *z-score* for each cluster. We defined the marker genes as genes with the highest correlation values, i.e. genes that represent the best the average pattern of the clusters. Keeping in mind that the marker genes should be easy to handle, we then selected the optimal marker genes displaying high expression levels while not belonging to extended protein families.

### Motif and transcription factor targets enrichment analysis

We performed enrichment analysis on the DEG in the different clusters for transcription factor targets genes and target motifs.

Motif discovery on the DEG set was performed using Find Individual Motif occurrences (FIMO) [36]. Motif list available for peach was obtained from PlantTFDB 4.0 [35]. To calculate the overrepresentation of motifs, DEGs were grouped by motif (grouping several genes and transcripts in which the motif was found). Overrepresentation of motifs was performed using hypergeometric tests using Hypergeometric {stats} available in R. Comparison was performed for the number of appearances of a motif in one cluster against the number of appearances on the overall set of DEG. As multiple testing implies the increment of false positives, *p-values* obtained were corrected using False Discovery Rate [96] correction method using p.adjust{stats} function available in R.

A list of predicted regulation between transcription factors and target genes is available for peach in PlantTFDB [35]. We collected the list and used it to analyse the overrepresentation of genes targeted by TF, using Hypergeometric {stats} available in R, comparing the number of appearances of a gene controlled by one TF in one cluster against the number of appearances on the overall set of DEG. *p-values* obtained were corrected using a false discovery rate as described above. Predicted gene homology to *Arabidopsis thaliana* and functions were retrieved from the data files available for *Prunus persica* (GDR, https://www.rosaceae.org/species/prunus_persica/genome_v2.0.a1).

### GO enrichment analysis

The list for the gene ontology (GO) terms was retrieved from the database resource PlantRegMap [35]. Using the topGO package [97], we performed an enrichment analysis on GO terms for biological processes, cellular components and molecular functions based on a classic Fisher algorithm. Enriched GO terms were filtered with a *p-value* < 0.005 and the ten GO terms with the lowest *p-value* were selected for representation.

### Marker genes selection and RT-qPCR analyses

The seven marker genes were selected based on the following criteria:

- Their expression presented the best correlation with the average expression profiles of their cluster.
- They were not members of large families (in order to reduce issues caused by redundancy).
- We only kept genes for which we could design high efficiency primers for RT-qPCR. Marker genes were not selected based on modelling fit, nor based on their function.

cDNA was synthetised from 1µg of total RNA using the iScript Reverse Transcriptase Kit (Bio-rad Cat no 1708891) in 20 µl of final volume. 2 µL of cDNA diluted to a third was used to perform the qPCR in a 20 µL total reaction volume. qPCRs were performed using a Roche LightCycler 480. Three biological replicates for each sample were performed. Primers used in this study for qPCR are: *PavCSLG3* F:CCAACCAACAAAGTTGACGA, R:CAACTCCCCCAAAAAGATGA; *PavMEE9*: F:CTGCAGCTGAACTGGAACAG, R:ACTCATCCATGGCACTCTCC; *PavSRP*: F:ACAGGATCTGGAAAGCCAAG, R:AGGGTGGCTCTGAAACACAG; *PavTCX2*: F:CTTCCCACAACGCCTTTACG, R:GGCTATGTCTCTCAAACTTGGA; *PavGH127*: F:GCCATTGGTTGTAGGGTTTG, R:ATCCCATTCAGCATTCGTTC; *PavUDP-GALT1* F:CAATGTTGCTGGAAACCTCA, R:GTTATTCCACATCCGACAGC; *PavPP2C* F:CTGTGCCTGAAGTGACACAGA, R:CTGCACTGCTTCTTGATTTG; *PavRPII* F:TGAAGCATACACCTATGATGATGAAG, R:CTTTGACAGCACCAGTAGATTCC; *PavEF1* F:CCCTTCGACTTCCACTTCAG, R:CACAAGCATACCAGGCTTCA. Primers were tested for non-specific products by separation on 1.5% agarose gel electrophoresis and by sequencing each amplicon. Realtime data were analyzed using custom R scripts. Expression was estimated for each gene in each sample using the relative standard curve method based on cDNA diluted standards. For the visualization of the marker genes’ relative expression, we normalized the RT-qPCR results for each marker gene by the average RT-qPCR data for the reference genes *PavRPII* and *PavEF1*.

### Bud stage predictive modelling

In order to predict the bud stage based on the marker genes transcriptomic data, we used TPM values for the marker genes to train and test several models. First, all samples were projected into a 2-dimensional space using PCA, to transform potentially correlated data to an orthogonal space. The new coordinates were used to train and test the models to predict the five bud stage categories. In addition, we tested the model on RT-qPCR data for samples harvested from the ‘Fertard’ cultivar. For the modelling purposes, expression data for the seven marker genes were normalized by the expression corresponding to the October sample. We chose the date of October as the reference because it corresponds to the beginning of dormancy and it was available for all cultivars. For each date, the October-normalized expression values of the seven marker genes were projected in the PCA 2-dimension plan calculated for the RNA-seq data and they were tested against the model trained on ‘Cristobalina’, ‘Garnet’ and ‘Regina’ RNA-seq data.

We tested five different models (Multinomial logistic regression, Random forest classifier, k-nearest neighbour classifier, multi-layer perceptron and support vector classification) for 500 different combination of training/testing RNA-seq datasets, all implemented using the scikit-learn Python package [98]. The models were 5-fold cross-validated to ensure the robustness of the coefficients and to reduce overfitting. The models accuracies were calculated as the percentage of correct predicted stages in the RNA-seq testing set and the RT-qPCR dataset. Results presented in Fig. S10 (Additional file 1) show that the highest model accuracies were obtained for the support vector classification and the multinomial logistic regression models. We selected the logistic regression model for this study because the coefficients are biologically relevant.

## LIST OF ABBREVIATIONS

ABA: abscisic acid
ABF2: ABSCISIC ACID RESPONSE ELEMENT-BINDING FACTOR 2
ABI5: ABSCISIC ACID INSENSITIVE 5
AG: AGAMOUS
AGL9: AGAMOUS-like 9
AGL20: AGAMOUS-like 20
AP3: APETALA3
AREB3: ABSCISIC ACID RESPONSE ELEMENT-BINDING PROTEIN 3
ATHB7: ARABIDOPSIS THALIANA HOMEOBOX 7
CBF/DREB: C-REPEAT/DRE BINDING FACTOR 2/DEHYDRATION RESPONSE ELEMENT-BINDING PROTEIN
CSLG3: Cellulose Synthase like G3
DAM: DORMANCY ASSOCIATED MADS-box
DEG: differentially expressed gene
DNA: desoxyribonucleic acid
EE: Evening element motif
EF1: Elongation factor 1
ERF: ethylene-responsive element
FD: FLOWERING LOCUS D
FIMO: Find Individual Motif occurrences
FLC: FLOWERING LOCUS C
GH127: Glycosyl Hydrolase 127
GPX6: GLUTATHION PEROXIDASE 6
GR: GLUTATHION REDUCTASE
GRF7: GROWTH-REGULATING FACTOR7
GST8: GLUTATHION S-TRANSFERASE8
GO: gene ontology
H3: Histone 3
LEA: LATE EMBRYOGENESIS ABUNDANT PROTEIN
LHY: LATE-ELONGATE HYPOCOTYL
LP1: LIPID TRANSFER PROTEIN1
MEE9: maternal effect embryo arrest 9
Padj: adjusted *p-value*
Pav: *Prunus avium*
PC: principal component
PCA: principal component analysis
PDCB3: PLASMODESMATA CALLOSE-BINDING PROTEIN 3
PIF4: PHYTOCHROME INTERACING FACTOR 4
PIL5: PHYTOCHROME INTERACING FACTOR 3 LIKE 5
PP2C: Phosphatase 2C
RNA: ribonucleic acid
RPII: ribonucleic acid polymerase II
RT-qPCR: quantitative reverse transcriptase polymerase chain reaction
RVE1/8: REVEILLE1/8
SEP3: SEPALLATA3
SPT: SPATULA
SRP: STRESS RESPONSIVE PROTEIN
TCX2: TESMIN/TSO1-like CXC 2
TF: transcription factor
TPM: transcripts per million reads
UDP-GalT1: UDP-Galactose transporter 1
ZTL: ZEITLUPE

## DECLARATIONS

### Ethics approval and consent to participate

Not applicable.

### Consent for Publication

Not applicable.

### Availability of data and materials

RNA-seq data that support the findings of this study have been deposited in the NCBI Gene Expression Omnibus under the accession code GSE130426.

The graphical web interface DorPatterns is available at the address: bwenden.shinyapps.io\DorPatterns.

Scripts and codes for data analysis and modelling will be available on github upon acceptation of the manuscript.

### Author contributions

SC, BW, ED and PAW designed the original research. MA and JCY participated to the project design. NV performed the RNA-seq and analysed the RNA-seq with SC and BW. MF performed the RT-qPCR. JAC performed the TF and motifs enrichment analysis. MT developed the model. NV, SC and BW wrote the article with the assistance of all the authors. All authors have read and approved the manuscript.

## Acknowledgments

We thank the Fruit Experimental Unit of INRA (Bordeaux-France) for growing and managing the trees, and Teresa Barreneche, Lydie Fouilhaux, Jacques Joly, Hélène Christman and Rémi Beauvieux for the help during the harvest and for the pictures. Many thanks to Dr Varodom Charoensawan (Mahidol University, Thailand) for providing scripts for mapping and gene expression count extraction.

## ADDITIONAL FILES

### Additional file 1

Figure S1 Field temperature during the sampling season

Figure S2 Separation of samples by dormancy stage using read counts for all genes

Figure S3 Enrichments in gene ontology terms in the ten clusters

Figure S4 Expression patterns for the transcription factors and their targets

Figure S5 Separation of samples by dormancy stage and cultivar using all genes

Figure S6 Analysis of the 100 genes that best contribute to the PCA dimensions 1 and 2

Figure S7 Clusters of expression patterns for differentially expressed genes in the sweet cherry cultivars ‘Regina’, ‘Cristobalina’ and ‘Garnet’

Figure S8 Separation of samples by dormancy stage and cultivar using the seven marker genes

Figure S9 Multinomial logistic regression model details

Figure S10 Comparison of the accuracy for the five tested models

### Additional file 2

Table S1 Description of the flower bud samples used for RNA-seq and RT-qPCR

Table S2 ’Garnet’ differentially expressed genes and their assigned clusters

Table S3 Transcription factors with enriched targets in the different clusters

Table S4 Enrichments in gene ontology terms in the transcription factors targets

Table S5 Transcription factors with targeted motif enrichment in the clusters

Table S6 Contribution of differentially expressed genes to the PCA dimensions 1 and 2

Table S7 RNA-seq mapped reads and gene count information

## REFERENCES

1. Heide OM, Prestrud AK. Low temperature, but not photoperiod, controls growth cessation and dormancy induction and release in apple and pear. Tree Physiol. 2005;25:109–14. http://www.ncbi.nlm.nih.gov/pubmed/15519992.

2. Allona I, Ramos A, Ibáñez C, Contreras A, Casado R, Aragoncillo C. Review. Molecular control of winter dormancy establishment in trees. Spanish J Agric Res. 2008;6:201–10.

3. Cooke JEK, Eriksson ME, Junttila O. The dynamic nature of bud dormancy in trees: environmental control and molecular mechanisms. Plant Cell Env. 2012;35:1707–28. doi:10.1111/j.1365-3040.2012.02552.x.

4. Maurya JP, Triozzi PM, Bhalerao RP, Perales M. Environmentally Sensitive Molecular Switches Drive Poplar Phenology. Front Plant Sci. 2018;9 December:1–8.

5. Olsen JE. Light and temperature sensing and signaling in induction of bud dormancy in woody plants. Plant Mol Biol. 2010;73:37–47.

6. Cline MG, Deppong DO. The role of apical dominance in paradormancy of temperate woody plants: A reappraisal. J Plant Physiol. 1999;155:350–6. doi:10.1016/S0176-1617(99)80116-3.

7. Lang G, Early J, Martin G, Darnell R. Endo-, para-, and ecodormancy: physiological terminology and classification for dormancy research. Hort Sci. 1987;22:371–7.

8. Considine MJ, Considine JA. On the language and physiology of dormancy and quiescence in plants. J Exp Bot. 2016;67:3189–203.

9. Badeck FW, Bondeau A, Böttcher K, Doktor D, Lucht W, Schaber JJ, et al. Responses of spring phenology to climate change. New Phytol. 2004;162:295–309. doi:10.1111/j.1469-8137.2004.01059.x.

10. Menzel A, Sparks TH, Estrella N, Koch E, Aasa A, Ahas R, et al. European phenological response to climate change matches the warming pattern. Glob Chang Biol. 2006;12:1969–76. doi:10.1111/j.1365-2486.2006.01193.x.

11. Vitasse Y, Lenz A, Körner C. The interaction between freezing tolerance and phenology in temperate deciduous trees. Front Plant Sci. 2014;5:541.

12. Bigler C, Bugmann H. Climate-induced shifts in leaf unfolding and frost risk of European trees and shrubs. Sci Rep. 2018;8:1–10. doi:10.1038/s41598-018-27893-1.

13. Fu YH, Zhao H, Piao S, Peaucelle M, Peng S, Zhou G, et al. Declining global warming effects on the phenology of spring leaf unfolding. Nature. 2015;526:104–7.

14. Legave J-M, Guédon Y, Malagi G, El Yaacoubi A, Bonhomme M. Differentiated Responses of Apple Tree Floral Phenology to Global Warming in Contrasting Climatic Regions. Front Plant Sci. 2015;6 December. doi:10.3389/fpls.2015.01054.

15. Erez A. Bud Dormancy; Phenomenon, Problems and Solutions in the Tropics and Subtropics. In: Temperate Fruit Crops in Warm Climates. 2000. p. 17–48.

16. Atkinson CJ, Brennan RM, Jones HG. Declining chilling and its impact on temperate perennial crops. Environ Exp Bot. 2013;91:48–62. doi:10.1016/j.envexpbot.2013.02.004.

17. Snyder RL, de Melo-abreu JP. Frost Protection : fundamentals, practice and economics. Rome; 2005.

18. Falavigna V da S, Guitton B, Costes E, Andrés F. I Want to (Bud) Break Free: The Potential Role of *DAM* and SVP-Like Genes in Regulating Dormancy Cycle in Temperate Fruit Trees. Front Plant Sci. 2019;9 January:1–17.

19. Zhong W, Gao Z, Zhuang W, Shi T, Zhang Z, Ni Z. Genome-wide expression profiles of seasonal bud dormancy at four critical stages in Japanese apricot. Plant Mol Biol. 2013;83:247–64. doi:10.1007/s11103-013-0086-4.

20. Khalil-Ur-Rehman M, Sun L, Li CX, Faheem M, Wang W, Tao JM. Comparative RNA-seq based transcriptomic analysis of bud dormancy in grape. BMC Plant Biol. 2017;17:1–11. doi:10.1186/s12870-016-0960-8.

21. Chao WS, Doğramacı M, Horvath DP, Anderson J V., Foley ME. Comparison of phytohormone levels and transcript profiles during seasonal dormancy transitions in underground adventitious buds of leafy spurge. Plant Mol Biol. 2017;94:281–302.

22. Zhang Z, Zhuo X, Zhao K, Zheng T, Han Y, Yuan C, et al. Transcriptome Profiles Reveal the Crucial Roles of Hormone and Sugar in the Bud Dormancy of Prunus mume. Sci Rep. 2018;8:1–15. doi:10.1038/s41598-018-23108-9.

23. Min Z, Zhao X, Li R, Yang B, Liu M, Fang Y. Comparative transcriptome analysis provides insight into differentially expressed genes related to bud dormancy in grapevine (Vitis vinifera). Sci Hortic (Amsterdam). 2017;225 March:213–20. doi:10.1016/j.scienta.2017.06.033.

24. Ueno S, Klopp C, Leplé JC, Derory J, Noirot C, Léger V, et al. Transcriptional profiling of bud dormancy induction and release in oak by next-generation sequencing. BMC Genomics. 2013;14:236.

25. Paul A, Jha A, Bhardwaj S, Singh S, Shankar R, Kumar S. RNA-seq-mediated transcriptome analysis of actively growing and winter dormant shoots identifies non-deciduous habit of evergreen tree tea during winters. Sci Rep. 2014;4:1–9.

26. Lesur I, Le Provost G, Bento P, Da Silva C, Leplé JC, Murat F, et al. The oak gene expression atlas: Insights into Fagaceae genome evolution and the discovery of genes regulated during bud dormancy release. BMC Genomics. 2015;16:112.

27. Takemura Y, Kuroki K, Shida Y, Araki S, Takeuchi Y, Tanaka K, et al. Comparative transcriptome analysis of the less-dormant taiwanese pear and the dormant Japanese pear during winter season. PLoS One. 2015;10.

28. Zhu Y, Li Y, Xin D, Chen W, Shao X, Wang Y, et al. RNA-Seq-based transcriptome analysis of dormant flower buds of Chinese cherry (Prunus pseudocerasus). Gene. 2015;555:362–76. doi:10.1016/j.gene.2014.11.032.

29. Kumar G, Rattan UK, Singh AK. Chilling-mediated DNA methylation changes during dormancy and its release reveal the importance of epigenetic regulation during winter dormancy in Apple (Malus x domestica Borkh.). PLoS One. 2016;11:1–25. doi:10.1371/journal.pone.0149934.

30. Beauvieux R, Wenden B, Dirlewanger E. Bud Dormancy in Perennial Fruit Tree Species : A Pivotal Role for Oxidative Cues. Front Plant Sci. 2018;9 May:1–13.

31. Lloret A, Badenes ML, Ríos G. Modulation of Dormancy and Growth Responses in Reproductive Buds of Temperate Trees. Front Plant Sci. 2018;9:1–12.

32. Campoy JA, Ruiz D, Egea J. Dormancy in temperate fruit trees in a global warming context: A review. Sci Hortic (Amsterdam). 2011;130:357–72. doi:10.1016/j.scienta.2011.07.011.

33. Wenden B, Campoy JA, Jensen M, López-Ortega G. Climatic Limiting Factors: Temperature. In: Quero-García J, Iezzoni A, Pulawska J, Lang G, editors. Cherries: Botany, Production and Uses. CABI Publishing; 2017. p. 166–88.

34. Heide OM. Interaction of photoperiod and temperature in the control of growth and dormancy of Prunus species. Sci Hortic (Amsterdam). 2008;115:309–14. doi:10.1016/j.scienta.2007.10.005.

35. Jin J, Tian F, Yang DC, Meng YQ, Kong L, Luo J, et al. PlantTFDB 4.0: Toward a central hub for transcription factors and regulatory interactions in plants. Nucleic Acids Res. 2017;45:D1040–5.

36. Grant CE, Bailey TL, Noble WS. FIMO: Scanning for occurrences of a given motif. Bioinformatics. 2011;27:1017–8.

37. Causier B, Schwarz-Sommer Z, Davies B. Floral organ identity: 20 years of ABCs. Semin Cell Dev Biol. 2010;21:73–9. doi:10.1016/j.semcdb.2009.10.005.

38. Lee S-J, Kang J-Y, Park H-J, Kim MD, Bae MS, Choi H, et al. DREB2C Interacts with ABF2, a bZIP Protein Regulating Abscisic Acid-Responsive Gene Expression, and Its Overexpression Affects Abscisic Acid Sensitivity. Plant Physiol. 2010;153:716–27.

39. Chen Y, Chen Z, Kang J, Kang D, Gu H, Qin G. *AtMYB14* Regulates Cold Tolerance in Arabidopsis. Plant Mol Biol Report. 2013;31:87–97.

40. Liu Q, Wang Z, Xu X, Zhang H, Li C. Genome-wide analysis of C2H2 zinc-finger family transcription factors and their responses to abiotic stresses in poplar (Populus trichocarpa). PLoS One. 2015;10:1–25.

41. Kiełbowicz-Matuk A. Involvement of plant C2H2-type zinc finger transcription factors in stress responses. Plant Sci. 2012;185–186:78–85.

42. Koornneef M, Léon-Kloosterziel KM, Schwartz SH, Zeevaart JAD. The genetic and molecular dissection of abscisic acid biosynthesis and signal transduction in Arabidopsis. Plant Physiol Biochem. 1998;36:83–9.

43. Doğramacı M, Horvath DP, Anderson J V. Dehydration-induced endodormancy in crown buds of leafy spurge highlights involvement of MAF3- and RVE1-like homologs, and hormone signaling cross-talk. Plant Mol Biol. 2014;86:409–24.

44. Franklin KA, Lee SH, Patel D, Kumar SV, Spartz AK, Gu C, et al. PHYTOCHROME-INTERACTING FACTOR 4 (PIF4) regulates auxin biosynthesis at high temperature. Proc Natl Acad Sci. 2011;108:20231–5.

45. Penfield S, Josse E-M, Kannangara R, Gilday AD, Halliday KJ, Graham IA. Cold and light control seed germination through the bHLH transcription factor SPATULA. Curr Biol. 2005;15:1998–2006.

46. Fadón E, Herrero M, Rodrigo J. Flower development in sweet cherry framed in the BBCH scale. Sci Hortic (Amsterdam). 2015;192:141–7. doi:10.1016/j.scienta.2015.05.027.

47. Geilen K, Böhmer M. Dynamic subnuclear relocalization of WRKY40, a potential new mechanism of ABA-dependent transcription factor regulation. Plant Signal Behav. 2015;10:e1106659.

48. Liu R, Xu Y-H, Jiang S-C, Lu K, Lu Y-F, Feng X-J, et al. Light-harvesting chlorophyll a/b-binding proteins, positively involved in abscisic acid signalling, require a transcription repressor, WRKY40, to balance their function. J Exp Bot. 2013;64:5443–56.

49. Pandey SP, Roccaro M, Schön M, Logemann E, Somssich IE. Transcriptional reprogramming regulated by WRKY18 and WRKY40 facilitates powdery mildew infection of Arabidopsis. Plant J. 2010;64:912–23.

50. Zhou J, Lee C, Zhong R, Ye Z-H. MYB58 and MYB63 Are Transcriptional Activators of the Lignin Biosynthetic Pathway during Secondary Cell Wall Formation in Arabidopsis. Plant Cell Online. 2009;21:248–66.

51. Gibbs DJ, Voß U, Harding SA, Fannon J, Moody LA, Yamada E, et al. AtMYB93 is a novel negative regulator of lateral root development in Arabidopsis. New Phytol. 2014;203:1194–207.

52. Simmons AR, Davies KA, Wang W, Liu Z, Bergmann DC. SOL1 and SOL2 regulate fate transition and cell divisions in the arabidopsis stomatal lineage. Dev. 2019;146.

53. Engin H, Ünal A. Examination of Flower Bud Initiation and Differentiation in Sweet Cherry and Peach by Scanning Electron Microscope. Turk J Agric. 2007;31:373–9.

54. Li Z, Reighard GL, Abbott AG, Bielenberg DG. Dormancy-associated MADS genes from the EVG locus of peach [Prunus persica (L.) Batsch] have distinct seasonal and photoperiodic expression patterns. J Exp Bot. 2009;60:3521–30. doi:10.1093/jxb/erp195.

55. Rothkegel K, Sánchez E, Montes C, Greve M, Tapia S, Bravo S, et al. DNA methylation and small interference RNAs participate in the regulation of MADS-box genes involved in dormancy in sweet cherry (Prunus avium L.). Tree Physiol. 2017; October:1–13.

56. Jiménez S, Reighard GL, Bielenberg DG. Gene expression of DAM5 and DAM6 is suppressed by chilling temperatures and inversely correlated with bud break rate. Plant Mol Biol. 2010;73:157– 67. doi:10.1007/s11103-010-9608-5.

57. Zhao K, Zhou Y, Ahmad S, Xu Z, Li Y, Yang W, et al. Comprehensive Cloning of Prunus mume Dormancy Associated MADS-Box Genes and Their Response in Flower Bud Development and Dormancy. Front Plant Sci. 2018;9 February:1–12. doi:10.3389/fpls.2018.00017.

58. Mimida N, Saito T, Moriguchi T, Suzuki A, Komori S, Wada M. Expression of DORMANCY-ASSOCIATED MADS-BOX (DAM)-like genes in apple. Biol Plant. 2015;59:237–44.

59. Doǧramaci M, Horvath DP, Chao WS, Foley ME, Christoffers MJ, Anderson J V. Low temperatures impact dormancy status, flowering competence, and transcript profiles in crown buds of leafy spurge. Plant Mol Biol. 2010;73:207–26.

60. Howe GT, Horvath DP, Dharmawardhana P, Priest HD, Mockler TC, Strauss SH. Extensive Transcriptome Changes During Natural Onset and Release of Vegetative Bud Dormancy in Populus. Front Plant Sci. 2015;6.

61. Fennell AY, Schlauch KA, Gouthu S, Deluc LG, Khadka V, Sreekantan L, et al. Short day transcriptomic programming during induction of dormancy in grapevine. Front Plant Sci. 2015;6:834.

62. Leida C, Conesa A, Llácer G, Badenes ML, Ríos G. Histone modifications and expression of DAM6 gene in peach are modulated during bud dormancy release in a cultivar-dependent manner. New Phytol. 2012;193:67–80. doi:10.1111/j.1469-8137.2011.03863.x.

63. Götz K-P, Chmielewski FM, Homann T, Huschek G, Matzneller P, Rawel HM. Seasonal changes of physiological parameters in sweet cherry (Prunus avium L.) buds. Sci Hortic (Amsterdam). 2014;172:183–90. doi:10.1016/j.scienta.2014.04.012.

64. Tuan PA, Bai S, Saito T, Ito A, Moriguchi T. Dormancy-Associated MADS-Box (DAM) and the Abscisic Acid Pathway Regulate Pear Endodormancy Through a Feedback Mechanism. Plant Cell Physiol. 2017;58:1378–90. doi:10.1093/pcp/pcx074.

65. Tylewicz S, Petterle A, Marttila S, Miskolczi P, Azeez A, Singh RK, et al. Photoperiodic control of seasonal growth is mediated by ABA acting on cell-cell communication. Science (80-). 2018;360:212–5.

66. Leida C, Conejero A, Arbona V, Gómez-Cadenas A, Llácer G, Badenes ML, et al. Chilling-dependent release of seed and bud dormancy in peach associates to common changes in gene expression. PLoS One. 2012;7:e35777. doi:10.1371/journal.pone.0035777.

67. Ruttink T, Arend M, Morreel K, Storme V, Rombauts S, Fromm J, et al. A molecular timetable for apical bud formation and dormancy induction in poplar. Plant Cell. 2007;19:2370–90. doi:10.1105/tpc.107.052811.

68. Lopez-Molina L, Mongrand S, McLachlin DT, Chait BT, Chua NH. ABI5 acts downstream of ABI3 to execute an ABA-dependent growth arrest during germination. Plant J. 2002;32:317–28.

69. Yamane H, Wada M, Honda C, Matsuura T, Ikeda Y, Hirayama T, et al. Overexpression of Prunus DAM6 inhibits growth, represses bud break competency of dormant buds and delays bud outgrowth in apple plants. PLoS One. 2019;14:1–24.

70. Singh RK, Miskolczi P, Maurya JP, Bhalerao RP. A Tree Ortholog of SHORT VEGETATIVE PHASE Floral Repressor Mediates Photoperiodic Control of Bud Dormancy. Curr Biol. 2019;29:128–133.e2. doi:10.1016/j.cub.2018.11.006.

71. Ibáñez C, Kozarewa I, Johansson M, Ogren E, Rohde A, Eriksson ME. Circadian clock components regulate entry and affect exit of seasonal dormancy as well as winter hardiness in Populus trees. Plant Physiol. 2010;153:1823–33. doi:10.1104/pp.110.158220.

72. Kozarewa I, Ibáñez C, Johansson M, Ogren E, Mozley D, Nylander E, et al. Alteration of PHYA expression change circadian rhythms and timing of bud set in Populus. Plant Mol Biol. 2010;73:143– 56. doi:10.1007/s11103-010-9619-2.

73. Ding J, Böhlenius H, Rühl MG, Chen P, Sane S, Zambrano JA, et al. GIGANTEA-like genes control seasonal growth cessation in Populus. New Phytol. 2018;218:1491–503.

74. Johansson M, Ramos-sánchez JM, Conde D, Ibáñez C, Takata N, Allona I, et al. Role of the Circadian Clock in Cold Acclimation and Winter Dormancy in Perennial Plants. In: Anderson J, editor. Advances in Plant Dormancy. Springer, Cham; 2015. p. 51–74.

75. Denardi Porto D, Bruneau M, Perini P, Anzanello R, Renou JJ-P, Santos HPD, et al. Transcription profiling of the chilling requirement for bud break in apples: a putative role for FLC-like genes. J Exp Bot. 2015;66:2659–72. doi:10.1093/jxb/erv061.

76. Meissner M, Orsini E, Ruschhaupt M, Melchinger AE, Hincha DK, Heyer AG. Mapping quantitative trait loci for freezing tolerance in a recombinant inbred line population of Arabidopsis thaliana accessions Tenela and C24 reveals REVEILLE1 as negative regulator of cold acclimation. Plant, Cell Environ. 2013;36:1256–67.

77. Jiang Z, Xu G, Jing Y, Tang W, Lin R. Phytochrome B and REVEILLE1/2-mediated signalling controls seed dormancy and germination in Arabidopsis. Nat Commun. 2016;7:1–10. doi:10.1038/ncomms12377.

78. Rawat R, Schwartz J, Jones MA, Sairanen I, Cheng Y, Andersson CR, et al. REVEILLE1, a Myb-like transcription factor, integrates the circadian clock and auxin pathways. Proc Natl Acad Sci. 2009;106:16883–8. doi:10.1073/pnas.0813035106.

79. Farinas B, Mas P. Histone acetylation and the circadian clock: A role for the MYB transcription factor RVE8/LCL5. Plant Signal Behav. 2011;6:541–3.

80. Pérez FJ, Vergara R, Rubio S. H2O2 is involved in the dormancy-breaking effect of hydrogen cyanamide in grapevine buds. Plant Growth Regul. 2008;55:149–55.

81. Vergara R, Rubio S, Pérez FJ. Hypoxia and hydrogen cyanamide induce bud-break and up-regulate hypoxic responsive genes (HRG) and VvFT in grapevine-buds. Plant Mol Biol. 2012;79:171–8.

82. Ophir R, Pang X, Halaly T, Venkateswari J, Lavee S, Galbraith D, et al. Gene-expression profiling of grape bud response to two alternative dormancy-release stimuli expose possible links between impaired mitochondrial activity, hypoxia, ethylene-ABA interplay and cell enlargement. Plant Mol Biol. 2009;71:403–23.

83. Horvath DP, Sung S, Kim D-H, Chao WS, Anderson J. Characterization, expression and function of DORMANCY ASSOCIATED MADS-BOX genes from leafy spurge. Plant Mol Biol. 2010;73:169–79.

84. de la Fuente L, Conesa A, Lloret A, Badenes ML, Ríos G. Genome-wide changes in histone H3 lysine 27 trimethylation associated with bud dormancy release in peach. Tree Genet Genomes. 2015;11:45. doi:10.1007/s11295-015-0869-7.

85. Saito T, Bai S, Imai T, Ito A, Nakajima I, Moriguchi T. Histone modification and signalling cascade of the dormancy-associated MADS-box gene, PpMADS13-1, in Japanese pear (Pyrus pyrifolia) during endodormancy. Plant Cell Environ. 2015;38:1157–66. doi:10.1111/pce.12469.

86. Pagnussat GC, Yu HJ, Ngo QA, Rajani S, Mayalagu S, Johnson CS, et al. Genetic and molecular identification of genes required for female gametophyte development and function in Arabidopsis. Development. 2005;132:603–14.

87. Rinne PL, van der Schoot C. Plasmodesmata at the crossroads between development, dormancy, and defense. Can J Bot. 2003;81:1182–97.

88. Marquat C, Vandamme M, Gendraud M, Pétel G. Dormancy in vegetative buds of peach: Relation between carbohydrate absorption potentials and carbohydrate concentration in the bud during dormancy and its release. Sci Hortic (Amsterdam). 1999;79:151–62.

89. Biswas S, Kerner K, Teixeira PJPL, Dangl JL, Jojic V, Wigge PA. Tradict enables accurate prediction of eukaryotic transcriptional states from 100 marker genes. Nat Commun. 2017;8:15309.

90. Meier U. Growth stages of mono-and dicotyledonous plants BBCH Monograph. 2001. http://pub.jki.bund.de/index.php/BBCH/article/view/515/464.

91. Bolger AM, Lohse M, Usadel B. Trimmomatic: A flexible trimmer for Illumina sequence data. Bioinformatics. 2014;30:2114–20.

92. Verde I, Jenkins J, Dondini L, Micali S, Pagliarani G, Vendramin E, et al. The Peach v2.0 release: high-resolution linkage mapping and deep resequencing improve chromosome-scale assembly and contiguity. BMC Genomics. 2017;18:225.

93. Trapnell C, Pachter L, Salzberg SL. TopHat: Discovering splice junctions with RNA-Seq. Bioinformatics. 2009;25:1105–11.

94. Wagner D. Chromatin regulation of plant development. Curr Opin Plant Biol. 2003;6:20–8.

95. Love MI, Huber W, Anders S. Moderated estimation of fold change and dispersion for RNA-seq data with DESeq2. Genome Biol. 2014;15:1–21.

96. Benjamini Y, Hochberg J. Controlling the false discovery rate: a practical and powerful approach to multiple testing. J R Stat Soc Ser B. 1995;57:289–300.

97. Alexa A, Rahnenführer J. topGO: Enrichment Analysis for Gene Ontology. R Packag version 2340. 2018. www.bioconductor.org.

98. Pedregosa F, Varoquaux G, Gramfort A, Michel V, Thirion B, Grisel O, et al. Scikit-learn: Machine Learning in Python. J Mach Learn Res. 2011;12:2825–30. doi:10.1007/s13398-014-0173-7.2.

