## Additional file 1 for "From bud formation to flowering: transcriptomic state defines the cherry developmental phases of sweet cherry bud dormancy"

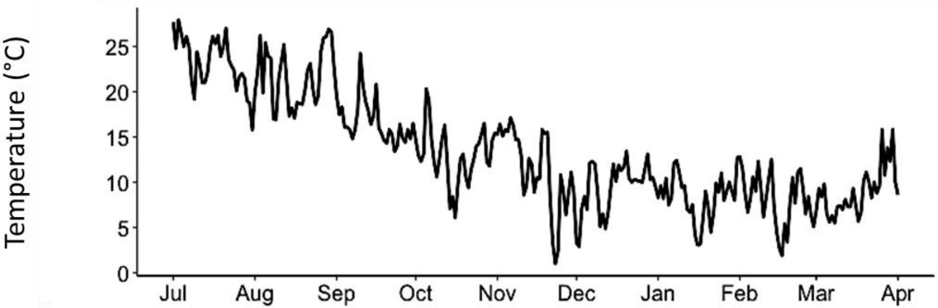


**Figure S1. Field temperature during the sampling season**

Average daily temperature recorded on site during the sampling season 2015/2016

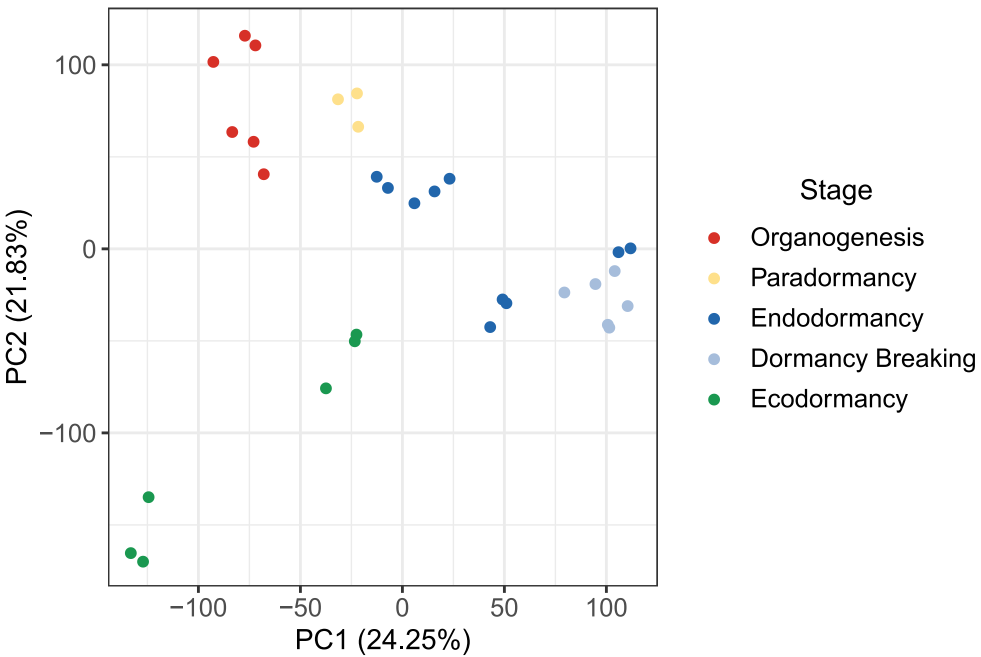


**Figure S2. Separation of samples by dormancy stage using read counts for all genes**.
The principal component analysis was conducted on the TPM (transcripts per million reads) values for the 26 873 genes in the flower buds of the cultivar 'Garnet', sampled on three trees between July 2015 and March 2016.

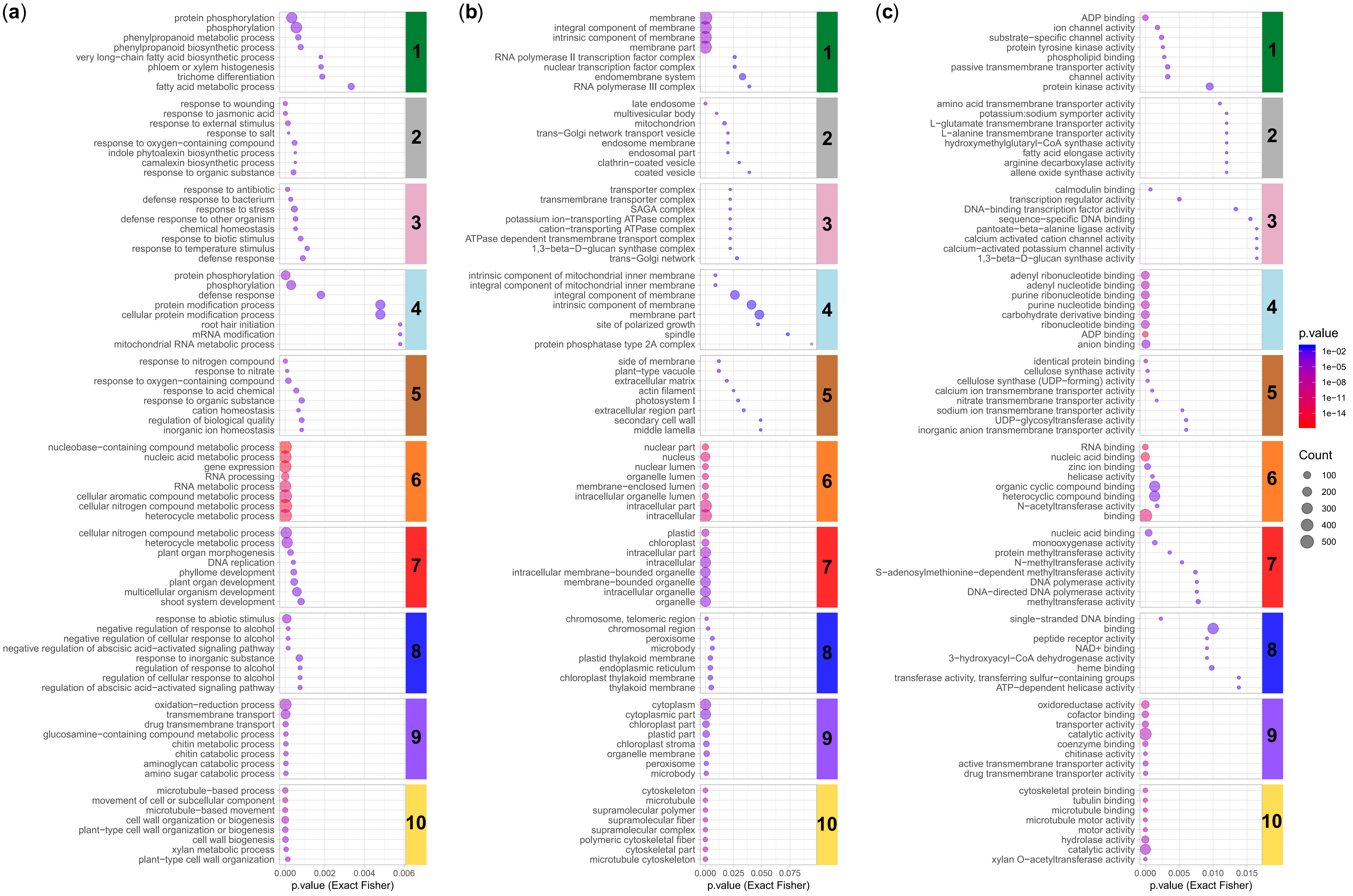


**Figure S3. Enrichments in gene ontology terms in the ten clusters**.

Using the topGO package (Alexa & Rahnenführer, 2018), we performed an enrichment analysis on GO terms for (a) biological processes, (b) cellular components and (c) molecular functions, based on a classic Fisher algorithm. Enriched GO terms with the lowest p-value were selected for representation. Dot size represents the number of genes belonging to the clusters associated with the GO term.

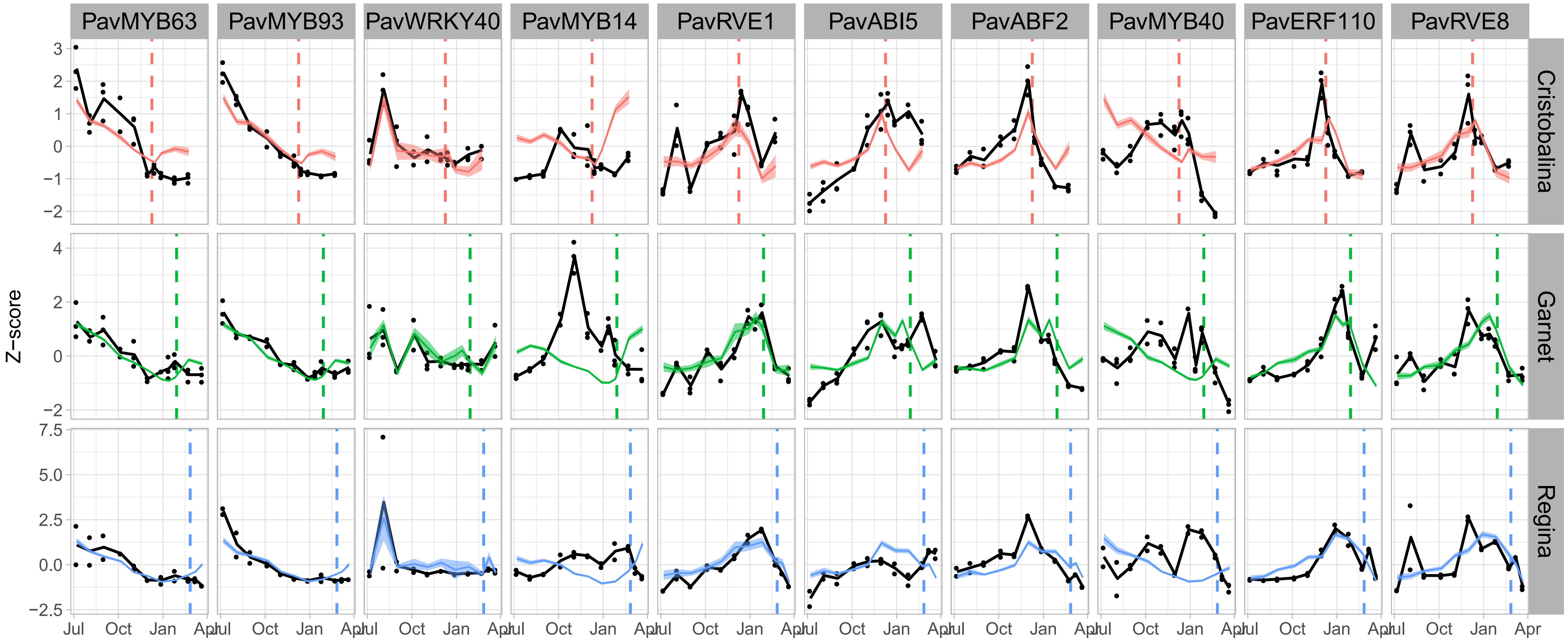


**Figure S4. Expression patterns for the transcription factors and their targets.**

Among the transcription factors identified with enriched targets in the different clusters (Table 1), ten belong to the differentially expressed genes set. Illustrated here are the z-scores calculated using the TPM for the DEGs: the transcription factor and the targets belonging to the clusters where the targets are enriched (see Table 1 for more details). Black lines represent the average z-score for the transcription factors for each cultivar. Colored lines and ribbons are the averaged z-scores and standard errors, respectively, for all differentially expressed target genes from the enriched cluster. Dash lines are the estimated date of dormancy release for each cultivar.

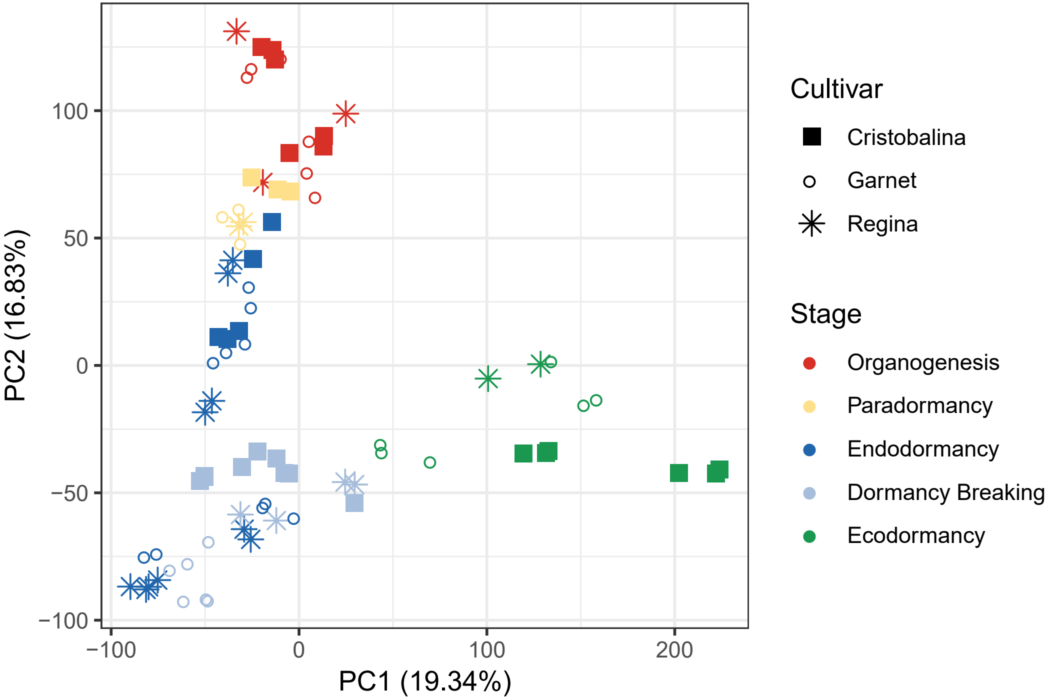


**Figure S5. Separation of samples by dormancy stage and cultivar using all genes**.
The principal component analysis was conducted on the TPM (transcripts per million reads) values for the 26 873 genes in the flower buds of the cultivars 'Cristobalina' (filled squares), 'Garnet' (empty circles) and 'Regina' (stars). Each point corresponds to one sampling time in a single tree.

(a)

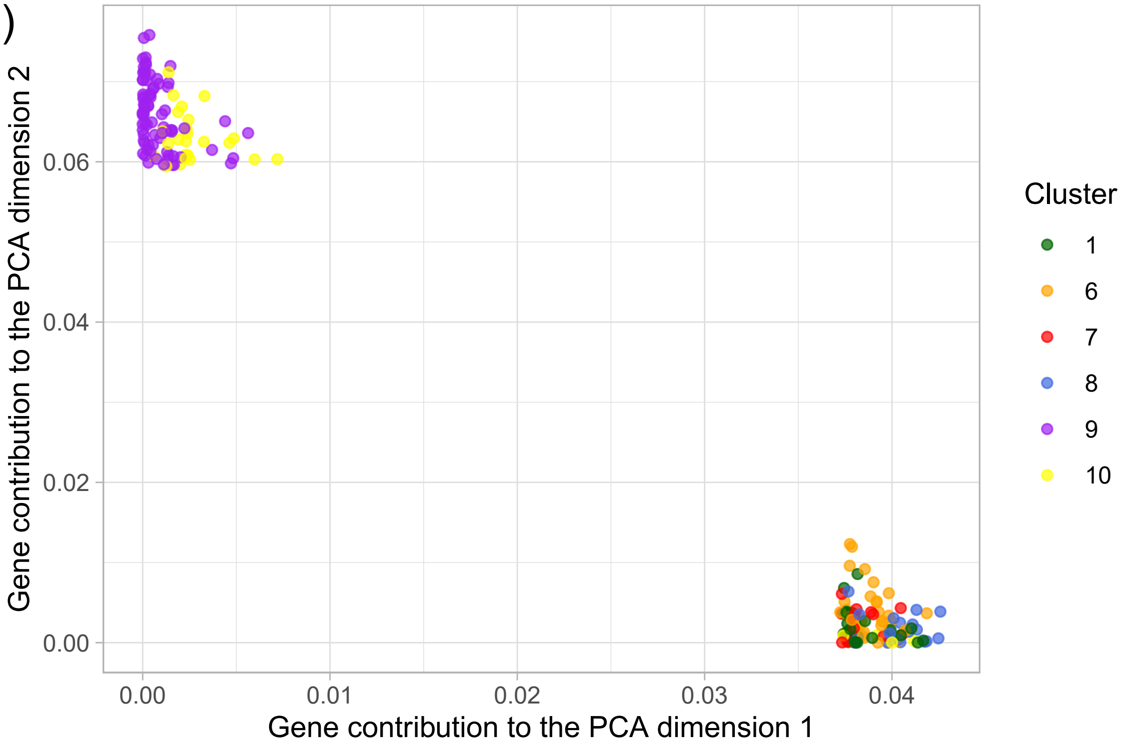


(b)

| PCA dimension | Cluster | Number of genes from the cluster in the 100 best contributors | p.value | adj. p.value | |
|---|---|---|---|---|---|
| Dim 1 | 1 | 22 | 0.55 | 0.69 | |
| | 6 | 36 | 2.94e-08 | 1.47e-07 | (***) |
| | 7 | 14 | 0.06 | 0.09 | |
| | 8 | 23 | 7.67e-06 | 1.92e-05 | (***) |
| | 10 | 5 | 0.97 | 0.97 | |
| Dim 2 | 9 | 76 | 4.48e-47 | 8.97e-47 | (***) |
| | 10 | 24 | 6.07e-05 | 6.07e-05 | (***) |

**Figure S6. Gene contribution to the PCA dimensions 1 and 2**

The principal component analysis was conducted on the TPM (transcripts per million reads) values for the 6883 differentially expressed genes (DEGs) in the flower buds of the cultivars 'Cristobalina', 'Garnet' and 'Regina'. (a) Contributions of the 100 best contributors of PCA dimensions 1 and 2. We assigned the cluster number from the cluster analysis performed on 'Garnet' DEGs. (b) Overrepresentation of the clusters in the 100 best contributors of PCA dimensions 1 and 2 was analysed using hypergeometric tests (Hypergeometric {stats} available in R). As multiple testing implies the increment of false positives, p-values were corrected using False Discovery Rate (Benjamini & Hochberg, 1995) correction method using p.adjust {stats} function available in R. (***): enrichment is significant with adjusted p-value < 0.005.

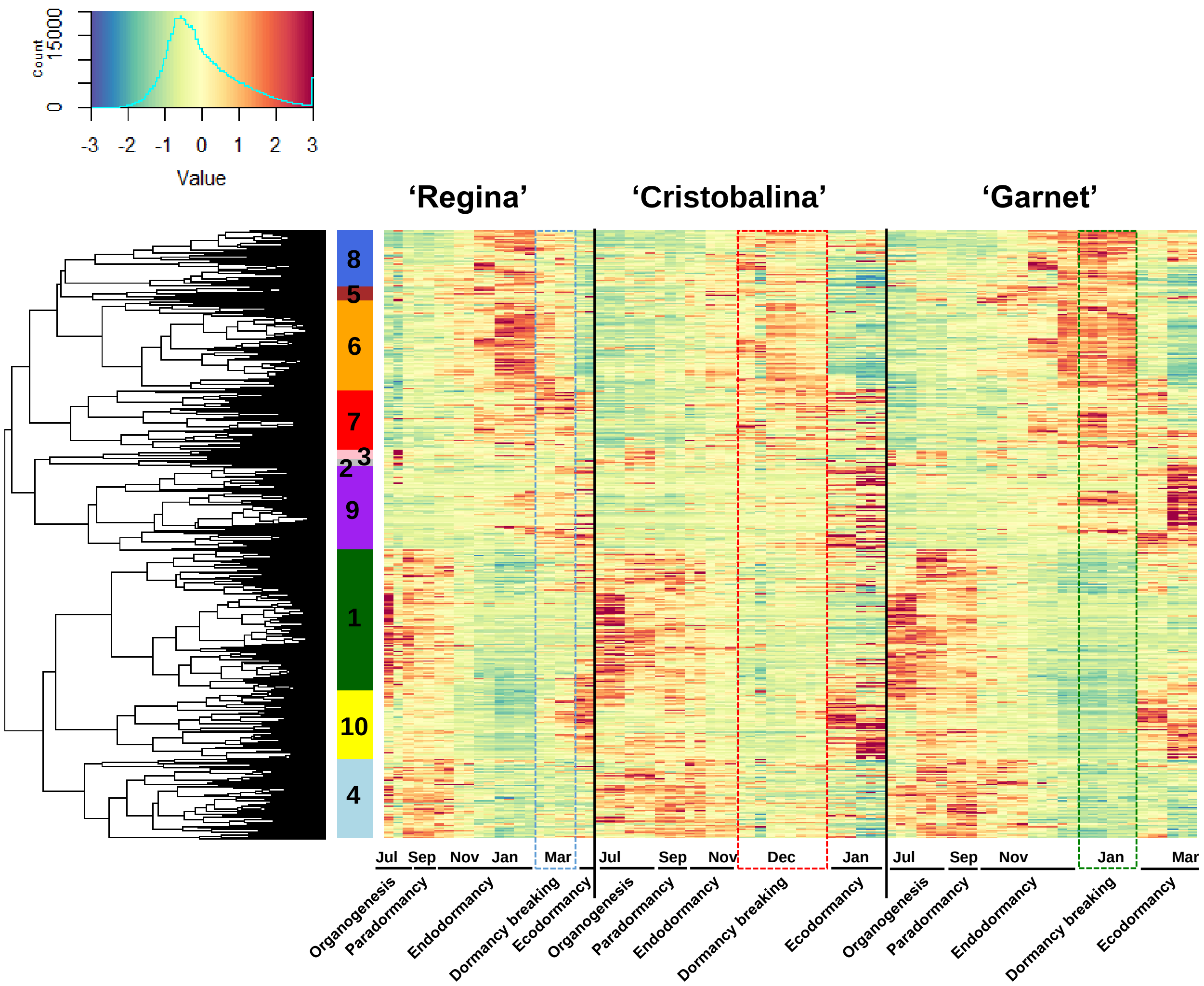


**Fig. S7 Clusters of expression patterns for differentially expressed genes in the sweet cherry cultivars 'Regina', 'Cristobalina' and 'Garnet'**
Heatmap for the 6 683 differentially expressed genes and corresponding clusters estimated using 'Garnet' data. Each column corresponds to the gene expression for flower buds from one single tree at a given date for a cultivar. Clusters are ordered based on the chronology of the expression peak (from earliest – July, 1-dark green cluster – to latest – March, 9 and 10). Expression values were normalized and *z-scores* are represented here.

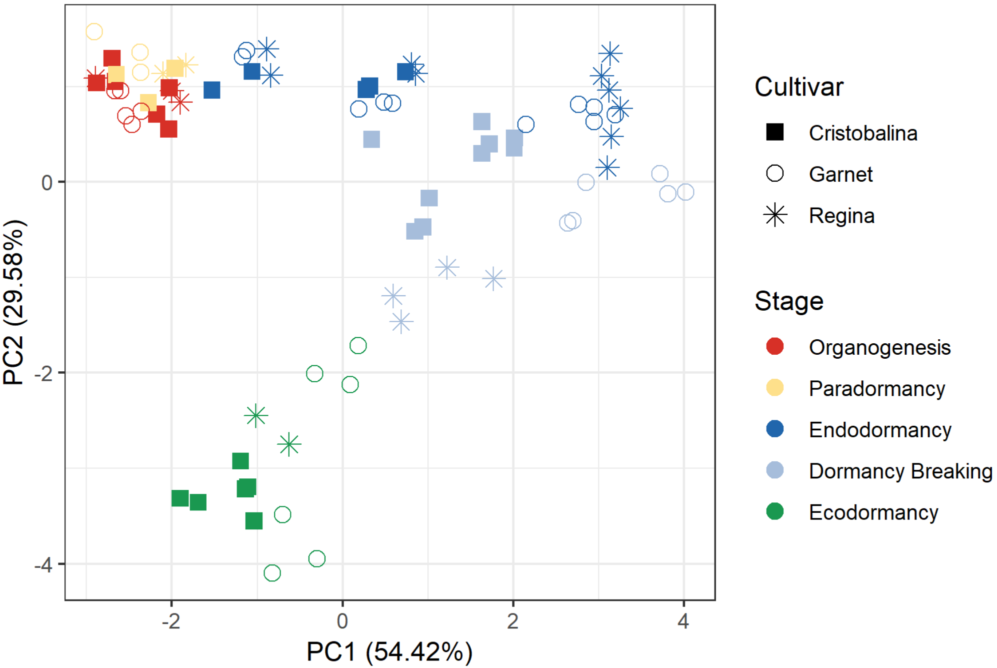


**Figure S8. Separation of samples by dormancy stage and cultivar using the seven marker genes**. The principal component analysis was conducted on the TPM (transcripts per million reads) values for the seven genes in the flower buds of the cultivars 'Cristobalina' (filled squares), 'Garnet' (empty circles) and 'Regina' (stars). Each point corresponds to one sampling time in a single tree.

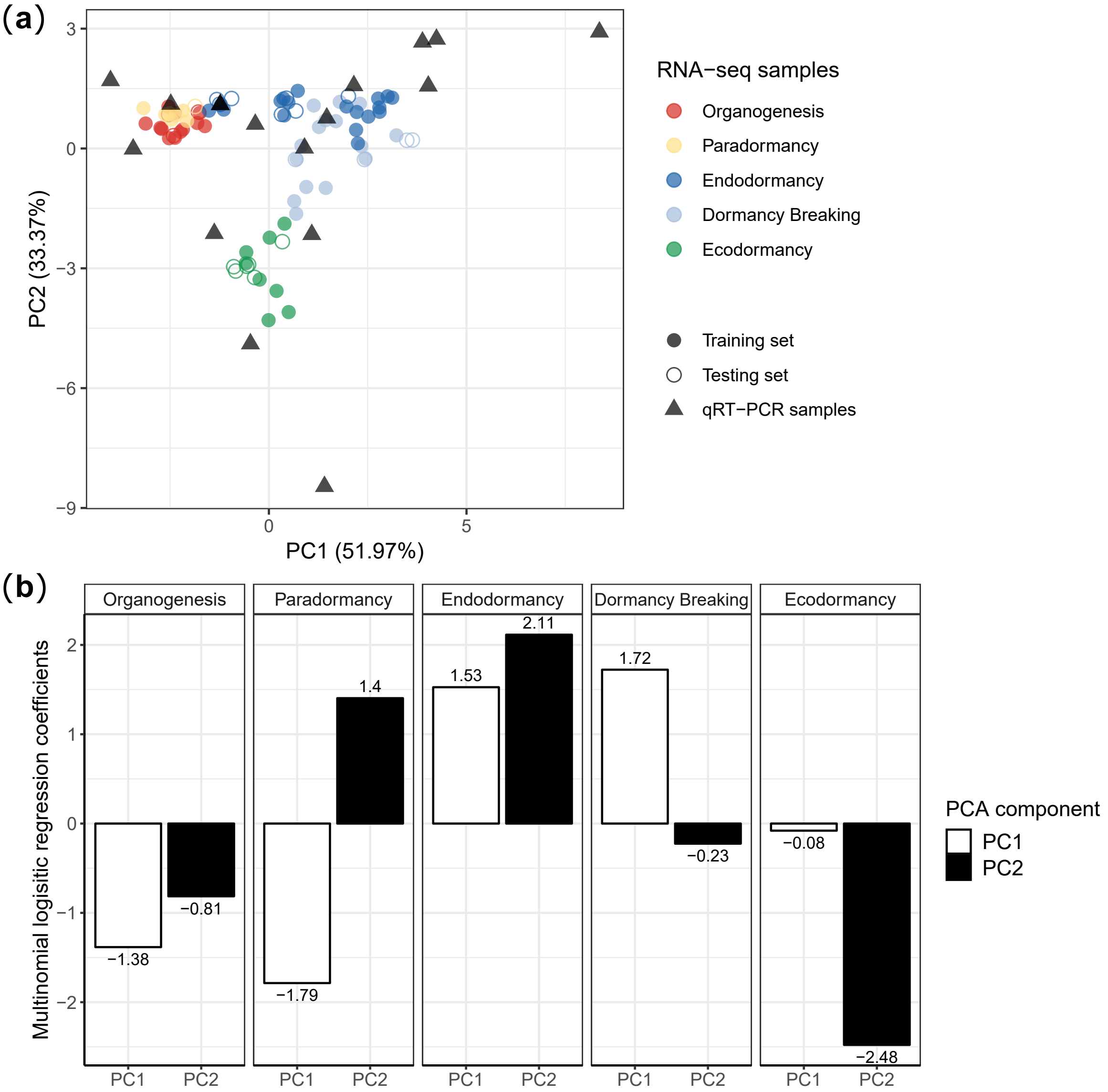


(c)

| Training Set | Testing Set | Model accuracy | | | Confusion Matrix – Reference Stages: Organogenesis | Paradormancy | Endodormancy | Dormancy Breaking | Ecodormancy |
|---|---|---|---|---|---|---|---|---|---|
| % RNA-seq data Randomly picked data subset (n = 60) | RNA-seq data Randomly picked data subset (n = 21) | 100% | Prediction | Organogenesis | 3 | 0 | 0 | 0 | 0 |
| | | | | Paradormancy | 0 | 1 | 0 | 0 | 0 |
| | | | | Endodormancy | 0 | 0 | 7 | 0 | 0 |
| | | | | Dormancy Breaking | 0 | 0 | 0 | 4 | 0 |
| | | | | Ecodormancy | 0 | 0 | 0 | 0 | 6 |
| | 'Fertard' qRT-PCR data (n = 17) | 71% | Prediction | Organogenesis | 1 | 0 | 0 | 0 | 0 |
| | | | | Paradormancy | 1 | 1 | 0 | 0 | 0 |
| | | | | Endodormancy | 0 | 0 | 9 | 0 | 0 |
| | | | | Dormancy Breaking | 0 | 0 | 2 | 0 | 0 |
| | | | | Ecodormancy | 0 | 0 | 1 | 1 | 1 |

**Figure S9. Multinomial logistic regression model details.**
(a) TPM (transcripts per million reads) values for the seven genes in the flower buds of the cultivars 'Cristobalina', 'Garnet' and 'Regina' were normalized by the october values, used as a reference and were then projected into a 2-dimensional space by PCA. Expression values for the seven marker genes were obtained for the late flowering cultivar 'Fertard' by qRT-PCR. These values were normalized by the October value and projected into the same PCA space. The model was then trained on the 2-D PCA projected data (filled circles) and tested on RNA-seq data (empty circles) and qRT-PCR data (filled triangles ). (b) Coefficients estimated for the multinomial logistic regression model that predicts the flower bud stages using the expression data projected into the 2-D dimansional space. (c) Prediction accuracy and confusion matrices for the model using RNA-seq and qRT-PCR testing subsets.

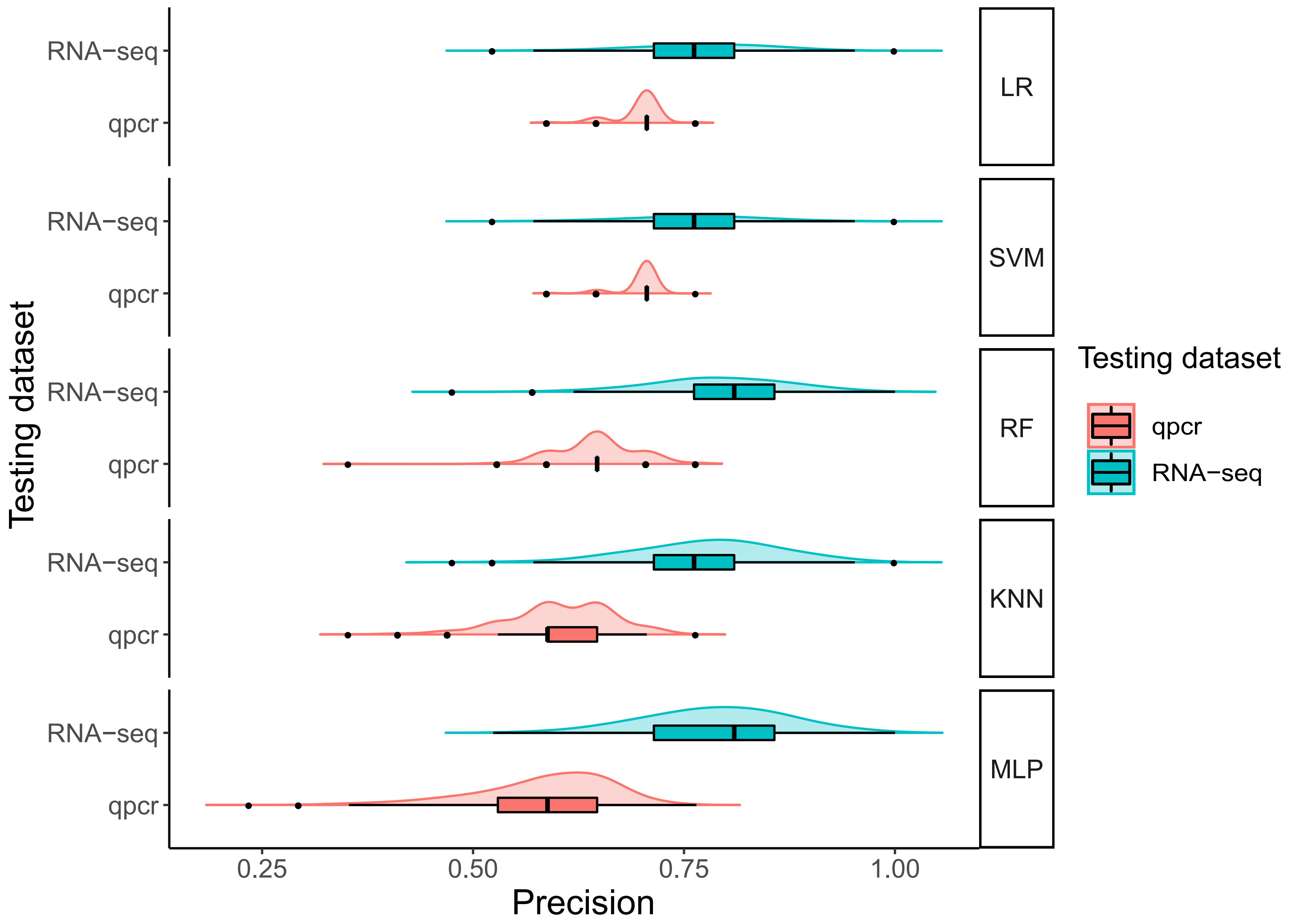


**Figure S10. Comparison of the accuracy for the five tested models**

For each model, we tested 500 random combination of RNA-seq data training and testing sets. For each training set, model parameters were estimated using 5 k-fold cross-validation and the best parameter sets were used to predict the bud development stages of the RNA-seq testing dataset and the qRT-PCR dataset. Precision is calculated as the percentage of correct stages predicted by the model. LR: logistic regression, SVM: support vector machines, RF: random forest, KNN: k-nearest neighbour, MLP: multi_layer perceptron.
